# Molecular forms of *Anopheles subpictus s.l.* and *Anopheles sundaicus s.l.* in the Indian subcontinent

**DOI:** 10.1101/2020.10.20.345272

**Authors:** Ankita Sindhania, Manoj K. Das, Gunjan Sharma, Sinnathamby N. Surendran, B.R. Kaushal, Himanshu P. Lohani, Om P. Singh

## Abstract

**Background:** *Anopheles subpictus s.l.* and *Anopheles sundaicus s.l.* are closely related species, each comprising of several sibling species. Ambiguities exist in the classification of these two nominal species and the specific status of members of *An. subpictus* complex. Identifying fixed molecular forms and mapping their spatial distribution will help in resolving the taxonomic ambiguities and understanding their relative epidemiological significance.

**Methods:** DNA sequencing of Internal Transcribed Spacer-2 (ITS2), 28S-rDNA (D1-to-D3 domains) and *cytochrome oxidase-II* of morphologically identified specimens of two nominal species, *An. subpictus s.l.* and *An. sundaicus s.l.* collected from the Indian subcontinent, was performed and subjected to genetic distance and molecular phylogenetic analyses.

**Results:** Molecular characterization of mosquitoes for rDNA revealed the presence of two molecular forms of *An. sundaicus s.l.* (identified as *An. epiroticus s.s.* and *An. sundaicus* D) and three molecular forms of *An. subpictus s.l.* (provisionally designated as Form A, B and C) in the Indian subcontinent. Phylogenetic analyses revealed two distinct clades: (i) subpictus clade, with a single molecular form of *An. subpictus* (Form A) prevalent in mainland India and Sri Lanka, and (ii) sundaicus clade, comprising of members of Sundaicus Complex., two molecular forms of *An. subpictus s.l.*, (Form B and C) prevalent in coastal areas or islands, and molecular forms reported from Thailand and Indonesia. Based on the number of float-ridges on eggs, all *An. subpictus* molecular Form B were classified as Species B whereas majority (80%) of the molecular Form A were classified as sibling species C. Fixed intragenomic sequence variation in ITS2 with the presence of two haplotypes was found in molecular Form A throughout its distribution.

**Conclusion:** A total of three molecular forms of *An. subpictus s.l.* and two molecular forms of *An. sundaicus s.l.* were recorded in the Indian subcontinent. Phylogenetically, two forms of *An. subpictus s.l.*, (Form B and C) prevalent in coastal areas or islands in the Indian subcontinent and molecular forms reported from Southeast Asia are members of Sundaicus Complex. Molecular Form A of *An. subpictus* is distantly related to all other forms and deserve a distinct specific status. Presence of *An. epiroticus* in Indian territory is recorded for the first time.

## Background

*Anopheles subpictus sensu lato* and *Anopheles sundaicus sensu lato* are closely related species belonging to Pyretophorus Series [1] and each has been reported to be comprised of several sibling members. *Anopheles subpictus s.l.* is widely distributed species prevalent throughout Oriental and Australasian Zones; mainly in Afghanistan, Australia, Bangladesh, Cambodia, China, India, Indonesia, Iran, Malaysia, Maldives, Mariana Islands, Myanmar (Burma), Nepal, New Guinea (Island)-Papua New Guinea, Pakistan, Philippines, Sri Lanka, Thailand and Vietnam [2]. In India, *An. subpictus s.l.* is the most common species occurring in all the mainland zones [3] and is found up to 1900 above msl and in many parts of Himalayas, though in small numbers. *Anopheles subpictus s.l.* gradually declines in abundance proceeding eastwards in India [4]. Other nominal taxa, *An. sundaicus s.l.*, has been recorded mainly from the coastal areas of north-eastern India, Andaman & Nicobar (A&N) Islands, Peninsular Malaysia, Malaysian Borneo (Miri, Sarawak), northern Sumatra & Java, and Indonesia [5]. Currently, the distribution of *An. sundaicus s.l.* in India is limited to the A&N Islands and Kutch (western coast of India) [6–7]. *Anopheles sundaicus s.l.* has been reported to be disappeared from Chilka (Odisha, India) [8], which has been reported earlier [9].

In India, *An. sundaicus s.l.* is considered as a potent malaria vector, but *An. Subpictus s.l.* has not been recognized yet as a malaria vector by the national malaria control programme [10]. However, *An. subpictus s.l.* is considered a potential vector in Southeast Asian countries [11,12,13,14]. In India, there are increasing evidences supporting the role of *An. subpictus* as a potential malaria vector, especially in coastal areas of India [15, 16] and Sri Lanka [17]. The varying vectorial potential in different geographical areas is most likely due to the presence of different biological species (cryptic species) present in *An. subpictus s.l*. In India, four sibling species of *An subpictus*, provisionally designated as species A, B, C and D have been recognized based on paracentric inversions present on the chromosome X [18]. However, the status of sibling species in other countries, except Sri Lanka (where species A and B were identified based on chromosomal inversions) [19], remains obscure. Though *An. subpictus s.l.* has been incriminated as a malaria vector in several places in India, mostly in coastal areas, the differential role of their sibling species in malaria transmission is not well understood. Apparently, species B is an efficient vector of malaria, which is prevalent in coastal areas of India [19] and Sri Lanka [20]. *Anopheles sundaicus s.l*., comprises of four sibling species [5], designated as *An. sundaicus s.s., An. epiroticus s.s.* Linton & Harbach (formerly, *An. sundaicus* species A) [21], *An. sundaicus* species D [22, 23] and *An. sundaicus* species E [24]. All of them act as predominant malaria vectors depending upon the location [25].

Morphologically, *An. subpictus s.l.* and *An. sundaicus s.l.* are almost similar and are distinguished based on the presence/absence of speckling on legs [26]. However, the taxonomic status of members of *An. subpictus* seems to be ambiguous based on the molecular phylogenetic studies which revealed that the majority of the morphologically attributed species B of *An. subpictus* is closely related to *An. sundaicus* complex and far distantly related to other members of *An. subpictus s.l.* [27–28]. Therefore, Surendran et al. (2010) [27] recognized species B of Subpictus Complex as a member of Sundaicus Complex. Interestingly, species B of Subpictus Complex [29] and members of the Sundaicus Complex are prevalent mainly in coastal areas and islands [14], although ecological plasticity in the breeding preference has also been noted [5].

Molecular characterization of *An. subpictus* from different geographical regions is important in identifying different biological forms, their distribution and role in malaria epidemiology. Studies carried out in Sri Lanka revealed the presence of two distinct molecular forms. Very recently, three additional molecular forms in respect of ITS2 has been reported from Thailand and Indonesia [30]. However, reliable published data on molecular forms of *An. subpictus* in Indian territory is not available. Few published and unpublished data from GenBank showed a high degree of polymorphism in ITS2 sequences, in the Indian population, to the extent that every individual was different [31–33], which raises suspicion in the quality of sequences. The verification of such data is warranted. Therefore, we attempted to characterize the rDNA of *An. subpictus s.l.* population from wide geographical areas in the Indian subcontinent in addition to closely related species *An. sundaicus s.l*.

For molecular characterization, we preferred nuclear ribosomal DNA (rDNA), particularly ITS2 and 28S rDNA, which have been extensively used in species identification due to their high uniformity in sequence in an interbreeding population, while differences exist between species [34]. Such uniformity of sequence in a species is believed to be due concerted evolution acting on rDNA, which tends to homogenize sequence [35]. The homogenization of rDNA is thought to be due to unequal crossing over of rDNA copies arranged in a tandem fashion, but the phenomenon has not been fully understood [36]. On the other hand mitochondrial DNA, *COI* in particular has also been successfully used in barcoding of life where sufficient gap, known as ‘barcoding gap’ (an arbitrary threshold), exists between intraspecific variation and interspecific divergence. In closely related species, such gap often doesn’t exist, but a high percentage of well-differentiated species has similar or even identical *COI* sequences [37], particularly in dipteran [38]. Such overlap may be due to the retention of ancestral polymorphism and introgression [39]. Moreover, rDNA is thought to be advantageous over the mtDNA in terms of the speed of lineage sorting of its multicopy rDNA array [39].

## Methodology

### Mosquito collection

Adult female *An. subpictus s.l.* and *An. sundaicus s.l.* mosquitoes were collected from different parts of the Indian subcontinent, mainly from mainland India, A&N Islands (India) and Sri Lanka. From mainland India, mosquitoes were collected from Ranchi (Jharkhand, 23°21’N, 85°18’E), Nuh (Haryana, 28°06’N, 76°59’E), Goa (15°38’N, 73°45’E), Alwar (Rajasthan, 27°37’N, 76°35’E), Jodhpur (Rajasthan, 26°18’N, 73°00’E), Gadchiroli (Maharashtra, 20°10’N, 80°00’E), Delhi (28°34’N, 77°01’E), villages surrounding Chilka lake (Odisha) and Puducherry. The villages surveyed surrounding Chilka lake, which is a large brackish water lagoon covering an area of over 1,100 km^2^ in Orrisa state, were Panasapada (19°43’N 85°31’E), Satapada (19°40’N 85°26’E), Brahmgiri (19°47’N 85°36’E), Sipakuda (19°23’N 85°04’E), Minsa (19°18’N 84°47’E), Siara (19°44’N 85°32’E) and Gambhari (19°72’N, 85°46’E). In Puducherry, mosquitoes were collected from Kaliankuppam (12°05’N 79°55’E), Munjalkuppam (12°02N 79°76’E), Sedarapet (12°00’N 79°44’E), Pillayarkuppam (11°48’N 79°46’E). From A&N Islands, mosquitoes were collected from Kimios, Car Nicobar (9°15’N 92°76’E) and Sippighat, Port Blair (11°59’N 92°68’E). In Sri Lanka, mosquitoes were collected from Kallady (7°42’N 81°42’E), Muthoor (8°27’N 81°16’E), and Chenkalady (7° 47 ‘N 81°35’ E). Few *An. sundaicus s.l.* mosquitoes collected from Thanbyuzayat Township, Mon State (15°58’N 97°44’E) were also included in the study. The geographical location of mosquito collection sites is depicted in **Figure 1**. Mosquitoes were either preserved in isopropanol or kept dried in microcentrifuge containing a piece of silica gel crystal. Where feasible, live female mosquitoes were transported to the laboratory for oviposition and identification of sibling species based on the number of float-ridges on eggs. The adult female mosquitoes were identified morphologically using keys by Christophers (1933) [26] and Reid (1968) [40].

**Figure 1.**
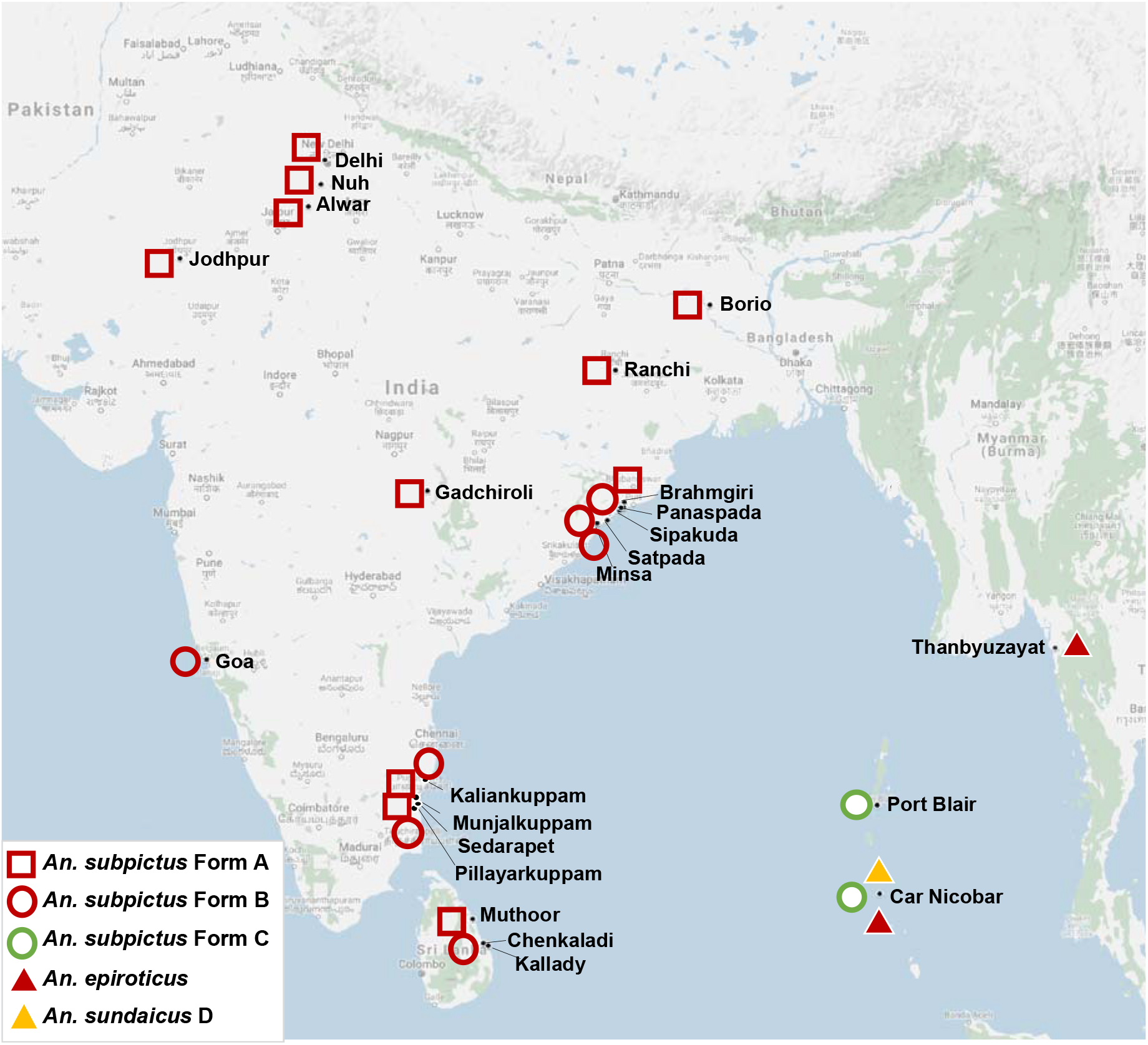
Map showing geographical locations of study sites and distribution of different molecular forms of *An. subpictus s.l.* and *An. sundaicus s.l.*

### Egg float-ridges count

Live *An. subpictus* mosquitoes were transported from mainland India only. The transported mosquitoes from the field were allowed to feed overnight on a-rabbit. Eggs were obtained from individual morphologically identified gravid-females in a plastic bowl with the inner side lined with blotting paper and containing a small amount of water for keeping the blotting paper moist. The mosquitoes, which successfully laid their egg, were persevered in a microcentrifuge tube containing isopropanol for molecular studies. Eggs from individual mosquitoes were transferred on to microslides and the number of ridges on egg-floats obtained were counted under a transmission microscope using 10X objective lens. At least 10 eggs from each mosquito were examined. Based on the mode number of ridges, the mosquitoes were classified for sibling species following Suguna et al. (1994) [18]

### DNA extraction, PCR amplification and sequencing

DNA was extracted from individual mosquitoes following Coen et al. (1982) [41].

Two adjacent regions of rDNA, i.e., ITS2 and partial 28S rDNA (D2-to-D3 domains), were PCR-amplified separately following Singh et al. (2006) [42]. In addition, to obtain a continuous stretch of rDNA covering partial 5.8S, full ITS2 and partial 28S rDNA (up to d3 domain) an approximately 2kb region was also amplified using primers ITS2A and D3B (**Table 1**) from at least two representative samples of each molecular form of *An*. *Subpictus s.l.* and *An. sundaicus s.l.* and one sample of *An. epiroticus*,. The PCR conditions were as described by Singh et al. (2006) [42], except extension time, which was increased to 1.5 minutes. The PCR products were cleaned using Exo-Sap IT (Thermo-Scientific) and sequenced using Big Dye Terminator v3.1 (Thermo-Scientific). The products were cleaned using the vendor’s protocol and electrophoresed in ABI-Prism 3770xl DNA sequencer. For the sequencing of ITS2 products, we used primers that were used for PCR amplification, i.e., ITS2A and ITS2B. Larger PCR products were sequenced using a primer walking strategy. The primers used for primer walking sequencing were ITS2A, ITS2B, D2A, D2B, D3A and D3B (sequences are provided in **Table 1**)

**Table 1.**
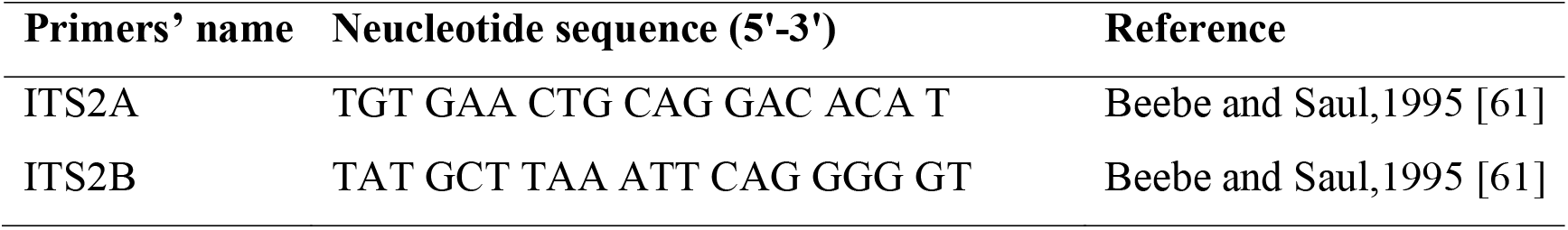
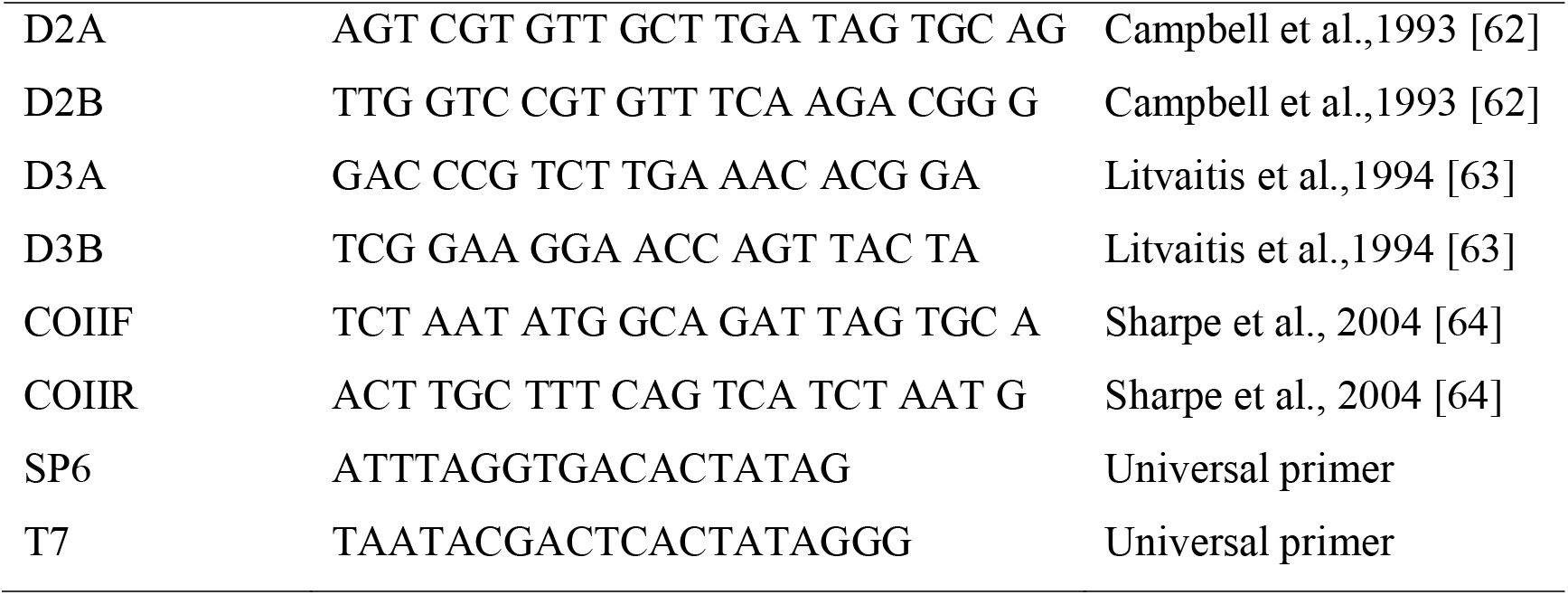
List of primers used for DNA sequencing

Five individuals of each molecular form, except *An. epiroticus* (n=1), were also sequenced for *cytochrome oxidase II* (*COII*) mitochondrial DNA. An approximately 750 bp region of the *COII* gene was PCR-amplified in a 20 μL reaction mixture using GOTaq Master Mix (Promega), 0.5 μM of each primer COIIF and COIIR (**Table 1**) and 0.5 μL of DNA. The PCR thermal cycling conditions were 95°C for 2 min followed by 35 cycles each at 95°C for 30 s, 55°C for 30 s, 72°C for 1 min and a final extension at 72°C for 7 min. The PCR products were cleaned using Exo-Sap IT (Thermo Scientific) and sequenced using Big Dye Terminator v3.1 (Thermo Scientific)..

### Cloning, sequencing and phasing of haplotypes

During the sequencing of ITS2 of a molecular form of *An. subpictus s.l.* (Form A) collected from mainland India and Sri Lanka, we noticed the presence of mixed bases in DNA sequence chromatogram starting from a specific point due to the presence of indel in one of the two haplotypes present in the mosquito (**Figure 2**). To phase out the sequence of two haplotypes, PCR product amplified for the ITS2 region of a total of five samples (three samples from mainland India, one each from Delhi, Ranchi and Kutch, and two samples from Sri Lanka) were sequenced after cloning. For cloning, briefly, PCR products were purified using Qiaquick PCR Purification Kit, cloned in pGEM-T Vector System (Promega) following vendor’s protocol and transformed into chemically competent *E. coli* DH5α. The transformants were plated on LB-Agar containing 100 μg/ml ampicillin. White colonies were picked up and plasmid DNA was isolated by boiling them in 50 μl of TE buffer. At least five clones from each mosquito were sequenced using universal primers T7 and SP6 (sequences provided in **Table 1**).

**Figure 2.**
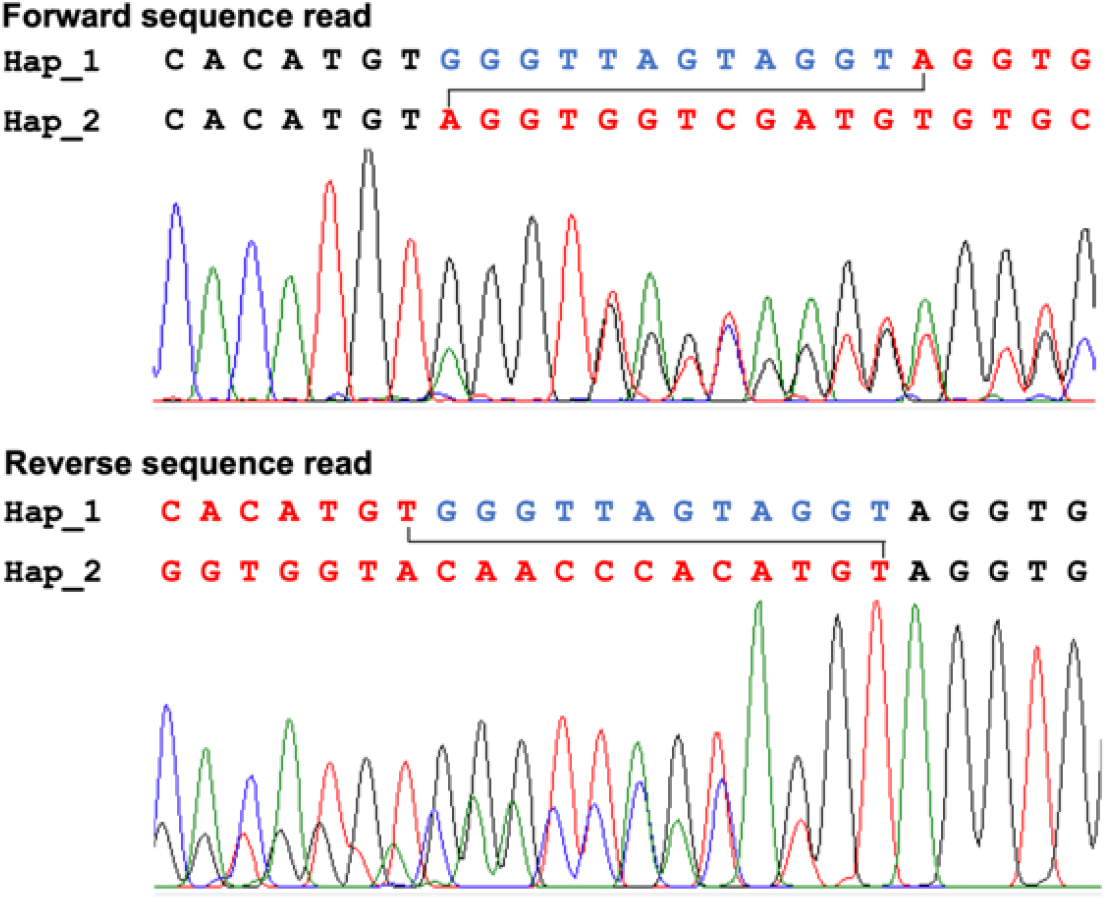
DNA sequence chromatograms of a portion of ITS2 (Forward and reverse) of *An. subpictus* form A, having indel in one haplotype. Hap_1 and Hap_2 in this figure represent haplotype 1 and 2, respectively.

### Sequence analyses and phylogenetic inferences

The DNA sequence chromatograms were edited using Finch TV (Geospiza, Inc.) and aligned using an online tool ClustalW 1.4 [43] available at https://www.genome.jp/tools-bin/clustalw. Since *An. subpictus* Form A had two haplotypes with 12 bp indel in ITS2, and sequence chromatograms were ambiguous from the point of indel, identification of two haplotypes were done on the basis of combining reads from forward sequences and reverse sequence (1X) up to the point of start of indel. The determination of sequence in the indel region was made empirically, as illustrated in **Figure 2**.

Phylogenetic analyses of ITS2 and 28S-D2-D3 were performed individually using software PAUP* version 4.0 beta 10 [44]. For phylogenetic analysis of ITS2, sequences of four *An. sundaicus s.l.* variants, i.e., *An. epiroticus s.s.* (variant I, GenBank accession no. AY768540), variant II (AY768541), *An. sundaicus* cytotype D. (variant III, AY768542), *An. sundaicus s.s.* (variant IV, AY768543) [24], *An. indefinitus* (GQ870332) and sequences of *An subpictus* variants from Thailand (MT068425 and MT068434) and Indonesia (MT068436) [30] were also used. Sequences of *An. fluviatilis* (DQ345964) [42] and *An. gambiae* (JN994138) were taken as outgroup taxa for ITS2 and *An. gambiae* (KC177663) and *An. fluvialitis* (DQ665846) [42] were taken as outgroup taxa for 28S-D2-D3 phylogenetic analysis. *Anopheles gambiae* was chosen as outgroup due to its phylogenetic proximity with *An*. *subpictus* and *An. sundaicus* complexes, whereas *An. fluviatilis* (belongs to Myzomyia series of subgenus *Cellia*) was chosen as it is phylogenetically relatively distant from *An. subpictus* and *An. sundaicus* complexes. We did not include ITS2 sequences of the Indian and Sri Lankan *An. subpictus* (published or submitted in GenBank) in the analysis due to significant error in the sequences (to the extent that every individual was different [31,32,33]), resulting from quality issues mostly arising due to the presence of indel in one of the two haplotypes present in molecular Form A, which is prevalent throughout mainland India and Sri Lanka. Maximum Likelihood (ML) methods were used for the construction of phylogenetic trees. The ML analyses were carried out using the heuristic searches and TBR branch-swapping algorithm with ten random taxon addition sequences. For ML-tree construction, the best evolutionary model of nucleotide substitution that fits the data were obtained using software Modeltest 3.7 [45]. The robustness of inference was calculated by the bootstrap method (1000 times) within PAUP. The estimated bootstrap values were reported as Maximum Likelihood bootstrap percentages (MLB).

### Allele-specific PCR (ASPCR) for the identification of molecular forms of *An. Subpictus* in mainland India

Since the sequencing of a large number of samples was not feasible, we developed a PCR assay for the quick identification of two molecular forms, *An. subpictus* Form A and Form B, only forms prevalent in mainland India. A universal primer Sub F (5’-ACT GCA GGA CAC ATG AAC ACC G-3’) was designed from upstream of ITS2A sequence and two reverse primers namely SUBP-A (5’-CGT TAC ACG CAA CAA GCG AC-3’) and SPB (5’-GCC GAC ACC ACC AAC TG-3’), specific to Form A and B of *An. subpictus*, respectively, were designed. The expected amplicon sizes for Form A and B of *An. subpictus* are 563 and 444 bp, respectively. The PCR reaction (15 μl) was carried out using GOTaq Master Mix (Promega) and 0.2 μM of each primer. The PCR thermal cycling conditions were 95°C for 2 min, followed by 35 cycles each at 95°C for 30 s, 58°C for 30 s, 72°C for 45 s, and a final extension at 72°C for 7 min. The PCR products were visualized on 2% agarose gel containing ethidium bromide under the gel documentation system. Numbers of samples on which ASPCR was performed have been given in **Table 2**.

**Table 2.** Number of mosquitoes identified into molecular forms of *An. subpictus* and An. sundaicus based on the rDNA (ITS2 and 28S-D2-D3) sequences and AS-PCR

| Locality | Molecular forms | DNA sequencing |  | AS-PCR* |
| --- | --- | --- | --- | --- |
|  |  | ITS2 | 28S-D2-D3 |  |
| Nuh, India | *An. subpictus* Form A | 4 | 4 | 86 |
| Ranchi, India | *An. subpictus* Form A | 63 | 19 | 15 |
| Borio, India | *An. subpictus* Form A | 2 | 2 | 10 |
| Alwar, India | *An. subpictus* Form A | 3 | 3 | 12 |
| Jodhpur, India | *An. subpictus* Form A | 2 | 7 | 7 |
| Gadchiroli, India | *An. subpictus* Form A | 4 | 4 | 25 |
| Chilka, India | *An. subpictus* Form A | 3 | 3 | 62 |
|  | *An. subpictus* Form B | 8 | 8 | 618 |
| Goa, India | *An. subpictus* Form B | 2 | 2 | 5 |
| Puducherry, India | *An. subpictus* Form A | 8 | 8 | 59 |
|  | *An. subpictus* Form B | 5 | 5 | 73 |
| Sri Lanka | *An. subpictus* Form A | 22 | 13 | 16 |
|  | *An. subpictus* Form B | 6 | 6 | 44 |
| Portblair, A&N Island | *An. subpictus* Form C | 5 | 5 | ND |
| Carnicobar, A&N Island | *An. subpictus* Form C | 5 | 5 | ND |
|  | *An. sondaicus* D (variant III#) | 12 | 12 | ND |
|  | *An. epiroticus* (variant I#) | 1 | 1 | ND |
| Myanmar | *An. epiroticus* (variant I#) | 4 | 4 | ND |

\* Assignment of Form B or C of *An. subpictus* in a specific locality was done on the basis of molecular form present in the area as determined by DNA sequencing.

## Results

### DNA sequences analyses

Details of samples sequenced for 28S-D2-D3 and ITS2 are provided in **Table 2**. Analysis of sequences revealed a total of five distinct molecular forms (in respect to ITS2 and 28S rDNA combined), of which three forms were from morphologically identified *An. subpictus s.l.* and two from *An. sundaicus s.l.* Sequences obtained from representative samples of each molecular forms (and haplotypes) after amplifying complete stretch of partial 5.8S rRNA, complete ITS2 and partial 28S rRNA (D1-D3 domain) have been shown in the aligned form in Figure S1. The sequences are also available from GenBank (accessions numbers: MW078484 to MW078490). The three forms of *An. subpictus* have been designated as molecular forms A, B and C. Form A of *An. subpictus s.l.* were recorded from mainland India and Sri Lanka, form B was found restricted to the coastal areas of mainland India (Chilka, Goa, Puducherry) and Sri Lanka island while form C from A&N Islands (Port Blair and Car Nicobar). The two molecular forms of *An. sundaicus s.l* were identified as *An. epiroticus s.s.* and *An. sundaicus* species D based on homology with published ITS2 sequences [21, 23, 24]. The characterization of species was done based on the polymorphism at three nucleotide positions of ITS2, i.e., 479 (T/C), 538 (G/T) and 603 (C/indel) (nucleotide position as described by Linton et al.(2005) [21]). Nucleotide bases at these three sites for *An. epiroticus, An. sundaicus* D and *An. sundaicus s.s.* were TGC, CGC and CT-(dash denotes indel), respectively. The *An. epiroticus s.s* was recorded from Myanmar and Car Nicobar (A&N Island) and *An. sundaicus* species D from Port Blair and Car Nicobar (A&N Island). The sequence of *An. epiroticus s.s.* recorded from Myanmar and A&N Islands had 100% homology to *An. epiroticus* s.s. [21] (variant I [24]). However, *An. epiroticus* from Car Nicobar was had mixed nucleotide bases (G+T) at the nucleotide position 538 (nucleotide position as per Linton et al.(2005)[21], where ‘G’ peak was substantially prominent as compared to ‘T’ (**Figure 3**). The *An. epiroticus* from Car Nicobar was identical to haplotype ‘TK’ designated by Zarowiecki et al. (2014) [58]. The ITS2 of *An. sundaicus* D was identical to variant III [24]. However, there was no difference in the 28S-D1-D3 sequence between *An. epiroticus* and *An. sundaicus* D. No individual variation in nucleotide sequences was found in each molecular form in a specific population.

**Figure 3.**
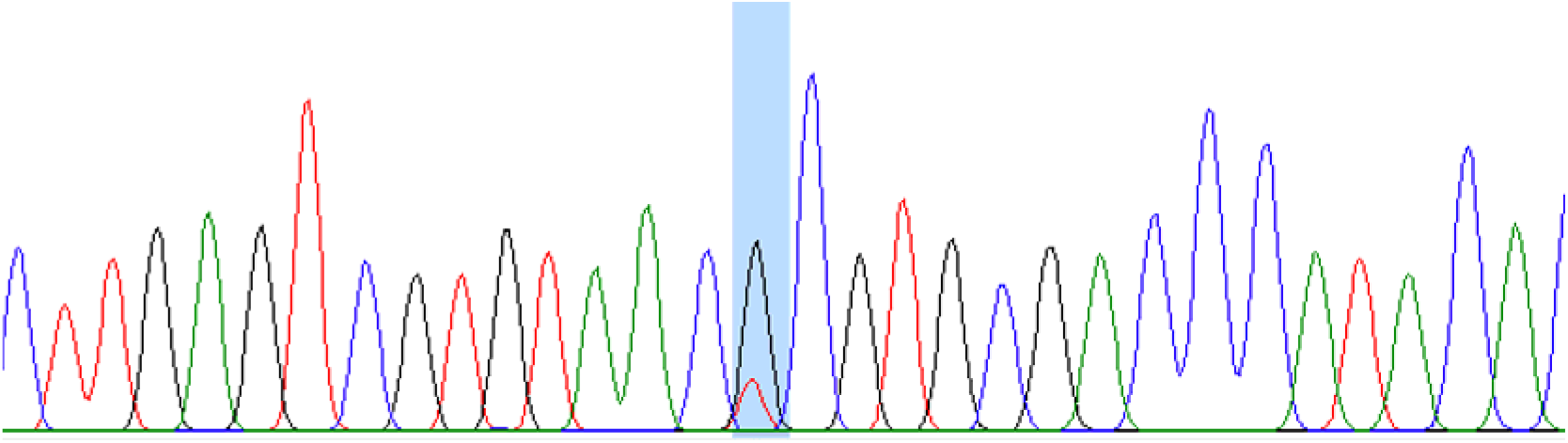
Portion of DNA chromatogram of ITS2 sequence of *An. epiroticus* from Car Nicobar island showing mixed bases (highlighted) at nucleotide base position 518 (base position as per Zarowiecki et al. 2014 [58])

### Intragenomic sequence variation in *An. subpictus* molecular Form A

DNA sequencing of ITS2 of *An. subpictus* (Form A) resulted in mixed nucleotide peaks from a specific position of nucleotide sequence (from base position 450 in Figure S1) resulting in an ambiguous sequence (**Figure 2**) from this point due to the presence of mixed haplotypes where one haplotype had indel. Phasing of haplotypes through sequencing of cloned PCR product revealed the presence of two haplotypes (referred to as haplotype 1 and haplotype 2 in Figure S1), which differ by indel/insert of 12. The presence of these two haplotypes was found fixed in Form A throughout its distribution from northern India to Sri Lanka.

### Genetic distance and phylogenetic analyses

The genetic distances and number of nucleotide substitutions, considering ITS2 and 28S-D2-D3 sequences, between three molecular forms of *An. subpictus* (A, B and C), *An. epiroticus* and *An. sundaicus* D are shown in **Table 3**. It was observed that *An. subpictus* Form A is distantly related to all other molecular forms of *An. subpictus s.l.* and *An. sundaicus s.l.* with nucleotide substitutions 100 to 106 and genetic distances 0.063 to 0.066 (Kimura-2-parameter). However, two molecular forms of *An. subpictus*, i.e., Form B and C, *An. epiroticus* and *An. sundaicus* species D are closely related (nucleotide substitution 1 to 10; Kimura-2-parameter genetic distance 0.001 to 0.006).

**Table 3.**
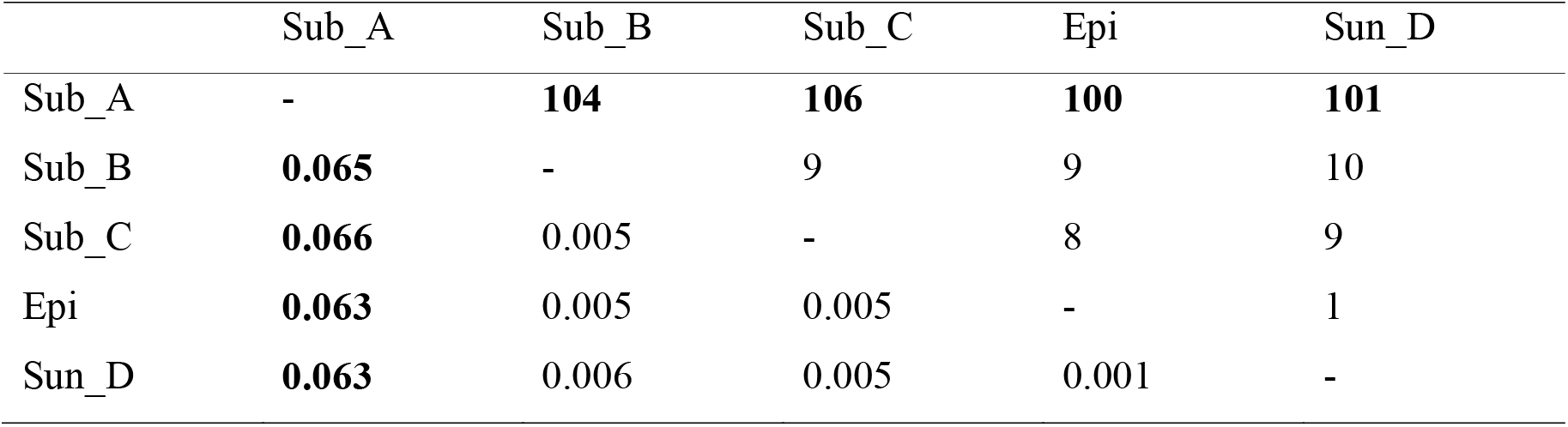
Pairwise distances (Kimura-2 parameter model) (below diagonal) and nucleotide substitution (above diagonal) between different molecular forms of *An. subpictus* and *An. sundaicus* based on ITS2 and 28S-D2-D3. Sub_A= *An. subpictus* Form A, Sub_B= *An. subpictus* Form B, Sub_C= *An. subpictus* Form C, Epi = *An. epiroticus* (variant I), Sun_D = *An. sundaicus* D (variant III)

Recently two new molecular forms of *An. subpictus s.l.* have been reported from Thailand and Indonesia with respect to ITS2 [30]. Therefore, a separate genetic distance analysis was performed, after including molecular forms reported from Thailand and Indonesia (representative GenBank sequences MT068425 and MT068436) [30] and *An. indefinitus* (GQ870332). The genetic distance and number of nucleotide substitutions are displayed in **Table 4**. It was observed that there is no genetic distance between Form C (India) and MT068436 (Indonesia) **Table 4**, but these sequences differed by 2 bp indel (Figure S1). *Anopheles subpictus* Forms B, C, molecular forms from Thailand and Indonesia (MT068425, MT068434, MT068436), *An. epiroticus s.s.*, *An. sundaicus* species D and *An. indefinitus* are closely related (nucleotide substitution 0 to 10; Kimura-2-parameter genetic distance 0.0 to 0.018) and are distantly related to Form A (nucleotide substitution 48 to 52; Kimura-2-parameter genetic distance 0.094 to 0.105).

**Table 4.** Nucleotide substitution (above diagonal) and pairwise distances (Kimura-2 parameter model) (below diagonal) and between different molecular forms of *An. subpictus* and *An. sundaicus* based on ITS2 sequences. Sub_A= *An. subpictus* Form A, Sub_B= *An. subpictus* Form B, MT068425 (Thailand), Sub_C= *An. subpictus* Form C, MT068436 (Indonesia), , Epi = *An. epiroticus* (variant I), Sun_D = *An. sundaicus* D (variant III), Ind=*An. indefinitus*

|  | Sub_A | Sub_B | MT068425 | Sub_C | MT068436 | Epi | Sun_D | Ind |
| --- | --- | --- | --- | --- | --- | --- | --- | --- |
| Sub_A | - | **51** | **49** | **52** | **52** | **47** | **48** | **48** |
| Sub_B | **0.103** | - | 2 | 4 | 4 | 5 | 6 | 3 |
| MT068425 | **0.099** | 0.004 | - | 4 | 4 | 3 | 4 | 1 |
| Sub_C | **0.105** | 0.008 | 0.008 | - | 0 | 7 | 8 | 5 |
| MT068436 | **0.105** | 0.008 | 0.008 | 0 | - | 7 | 8 | 5 |
| Epi | **0.094** | 0.010 | 0.006 | 0.013 | 0.013 | - | 1 | 4 |
| Sun_D | **0.097** | 0.011 | 0.008 | 0.015 | 0.015 | 0.002 | - | 5 |
| Ind | **0.097** | 0.006 | 0.002 | 0.009 | 0.009 | 0.008 | 0.010 | - |

The phylogenetic analysis of all molecular forms of *An. subpictus* and *An. Sundaicus s.l.* from the Indian subcontinent alongwith four variants in *An. sundaicus s.l.*, *An. indefinitus* and *An. subpictus s.l.* variants from Thailand and Indonesia were performed using ITS2 sequences. The best-fit model inferred for ITS2 sequences with outgroup taxa TPM2uf+G was taken for maximum likelihood (ML) analysis. Construction of phylogenetic tree resulted in two distinct monophyletic clades supported by high bootstrap value (100), i.e., subpictus clade comprising of the single molecular form of *An. subpictus* (Form A) and sundaicus clade comprising of *An. subpictus* Form B, *An. subpictus* Form C, molecular forms of *An. subpictus* from Thailand and Indonesia, all variants of *An. sundaicus s.l.* and *An. indefinitus* (**Figure 4**).

**Figure 4.**
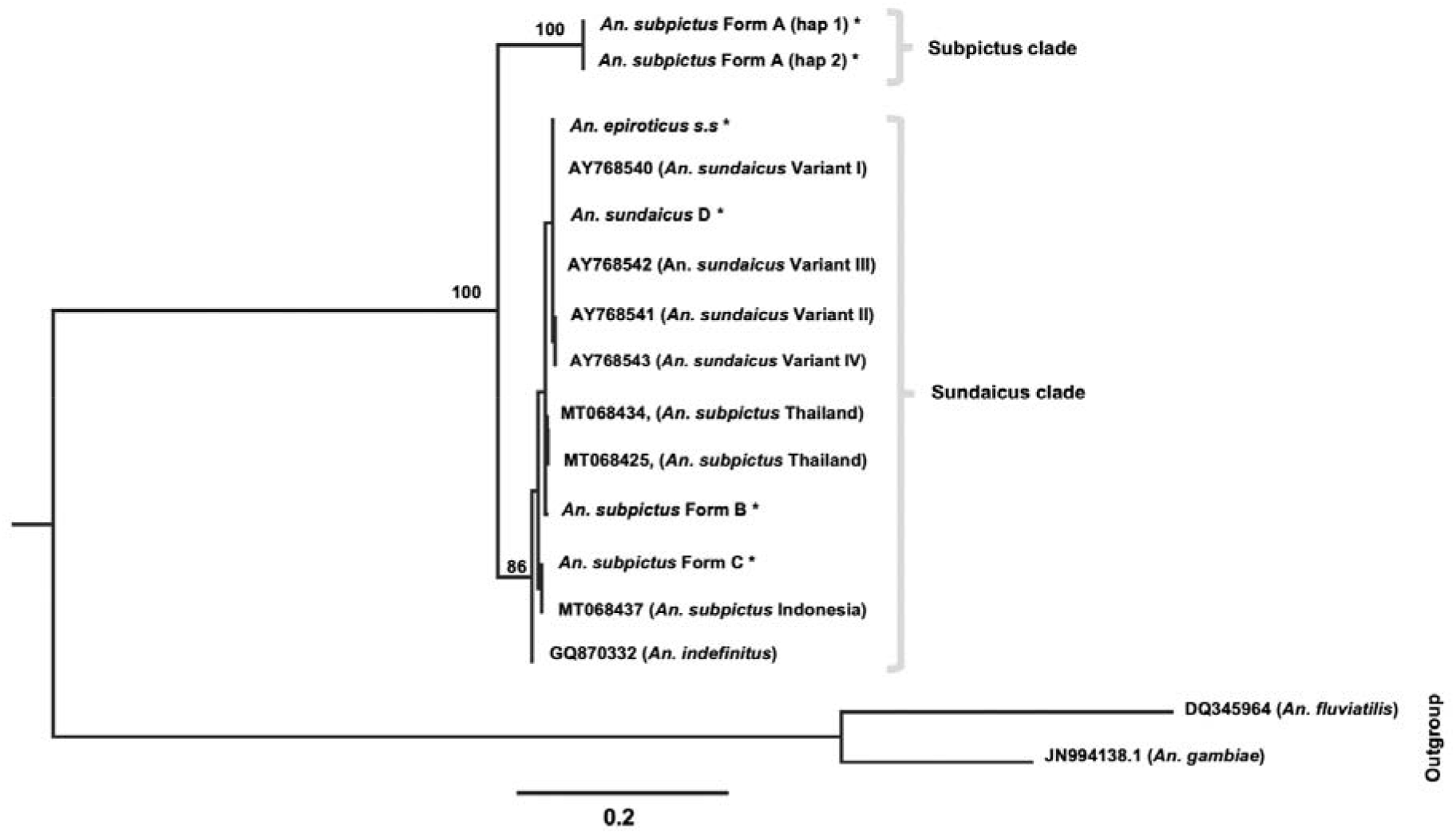
Maximum Likelihood (ML) rooted tree of different molecular forms of *An. subpictus* and *An. sundaicus* complex based upon ITS2 sequence data. Numbers at nodes are ML bootstrap values (only > 80 are shown). The sequence name with astricks (*) are the sequences generated in this study.

Phylogenetic tree was also constructed for 28S rDNA (D2-D3) sequences of all molecular forms of *An. subpictus* (Forms A, B and C), *An. sundaicus* D and *An. epiroticus s.s.* prevalent in the Indian subcontinent. The best-fit model of nucleotide substitution TIM3+G was implemented in the construction of ML trees. Cladogram derived from 28S rDNA sequences also resulted in the generation of two distinct monophyletic clades similar to ITS2, supported with high bootstrap value (100) (**Figure 5**).

**Figure 5.**
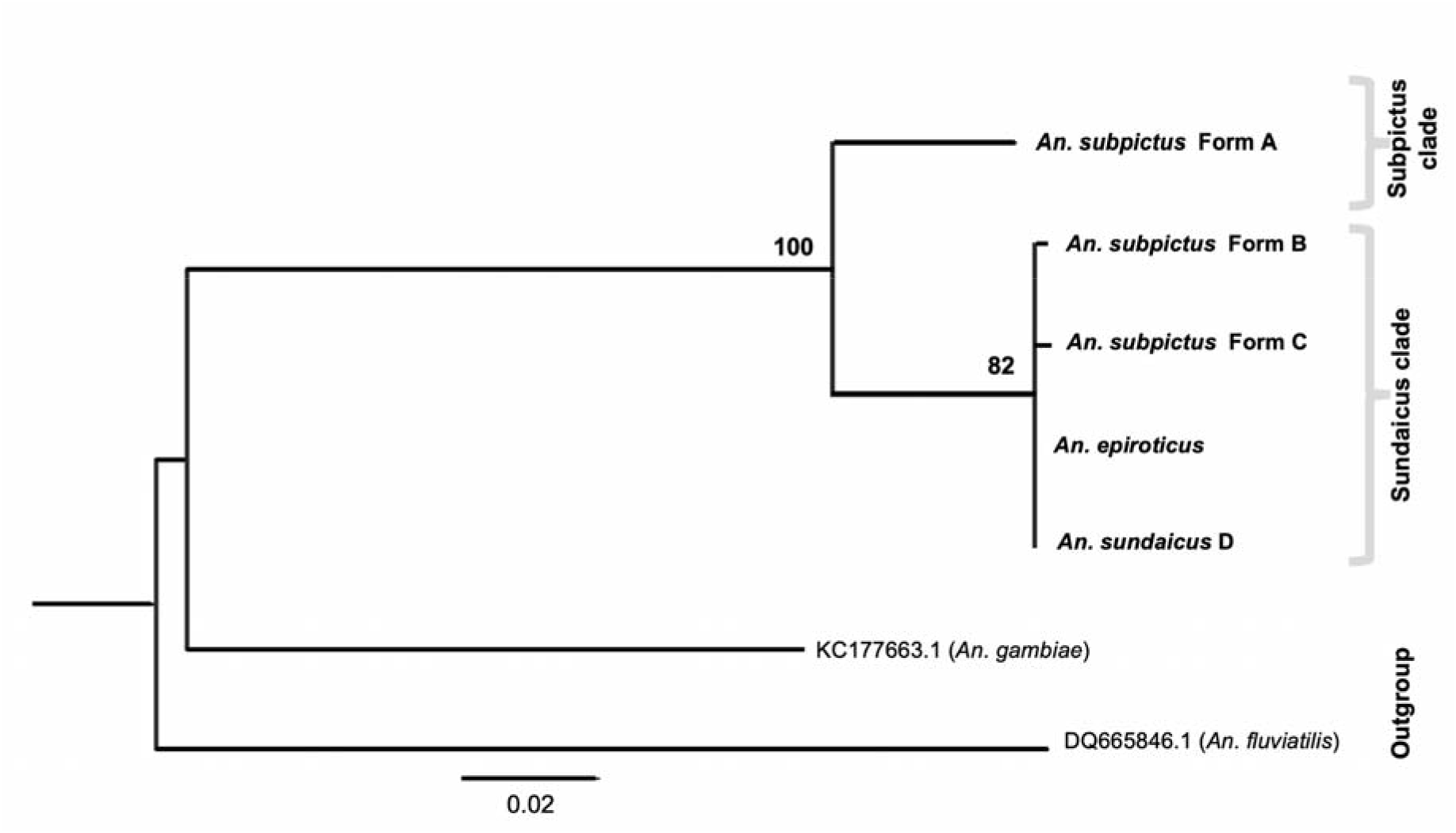
Maximum Likelihood (ML) rooted tree of different molecular forms of *An. subpictus* and *An. sundaicus* complex based upon 28S-D2-D3 sequences. Numbers at nodes are ML bootstrap values (only > 80 are shown).

Analysis of limited numbers of *COII* sequences of all representative molecular forms based on rDNA sequences revealed the presence of multiple haplotypes in each form. We recorded 3/5 (number of haplotypes/number sequenced) haplotypes in *An. subpictus* Form A, 3/5 in Form B, 1/5 in Form C, 2/5 in *An. sundaicus* D and 1/1 in *An. epiroticus.* The Maximum Likelihood phylogeny estimate based on *COII* data for all molecular forms of Indian *An. subpictus* and *An. sundaicus* complex was constructed using software PAUP 4.0 beta. The best fit model used was GTR+I+G, as estimated by the Modeltest 3.7 [45] (**Figure 6**). ML bootstrap values based on 1000 replicates were calculated. ML tree generated two distinct monophyletic clades similar to the phylogenetic tree based on rDNA. Based on *COII* phylogeny, *An. subpictus* form C was much closer to *An. epiroticus* than any members of *An. subpictus*.

**Figure 6.**
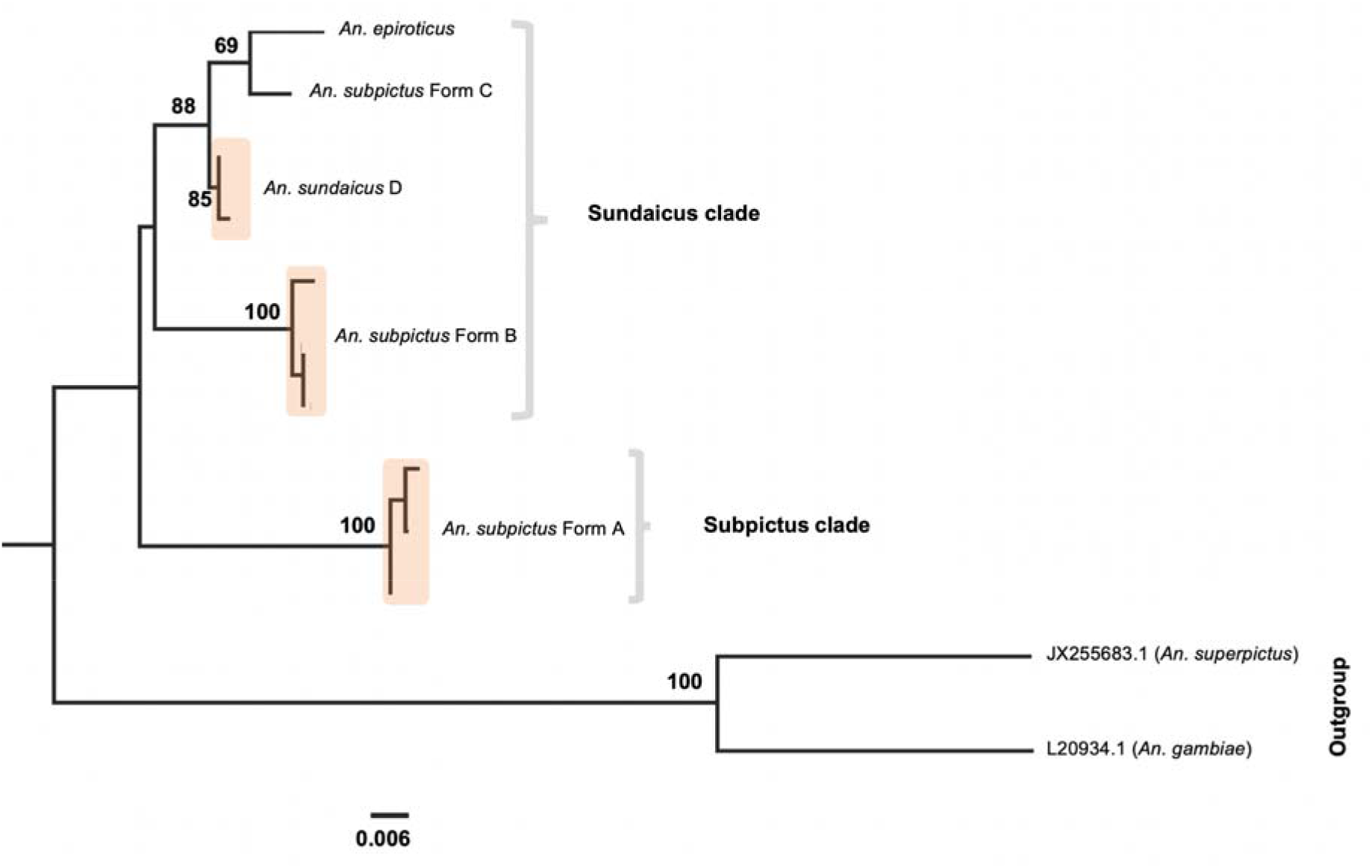
Maximum Likelihood (ML) rooted tree of different molecular forms of *An. subpictus* and *An. sundaicus* complex based upon *COII* sequences. Numbers at nodes are ML bootstrap values (only >50 are shown).

### Allele-specific PCR

Though this PCR was developed with an intension to differentiate the two forms of morphologically identified *An. subpictus s.l.* (*An. subpictus* Form A and Form B) in the Indian mainland and Sri Lanka, but upon discovery of additional species under the sundaicus clade (i.e., *An. subpictus* Form C) from A&N Islands, we found that this PCR cannot differentiate *An. subpictus* Form B from *An. subpictus* Form C. However, this PCR can discriminate members of subpictus clade (*An. subpictus* A) and sundaicus clade (*An. subpictus* Form B, C, *An. sundaicus* D and *An. epiroticus*) (**Figure 7**). This PCR was therefore used to differentiate molecular forms A and B of *An. subpictus s.l.* in areas where Form C or *An. sundaicus s.l.* was absent (mainland India and Sri Lanka).

**Figure 7.**
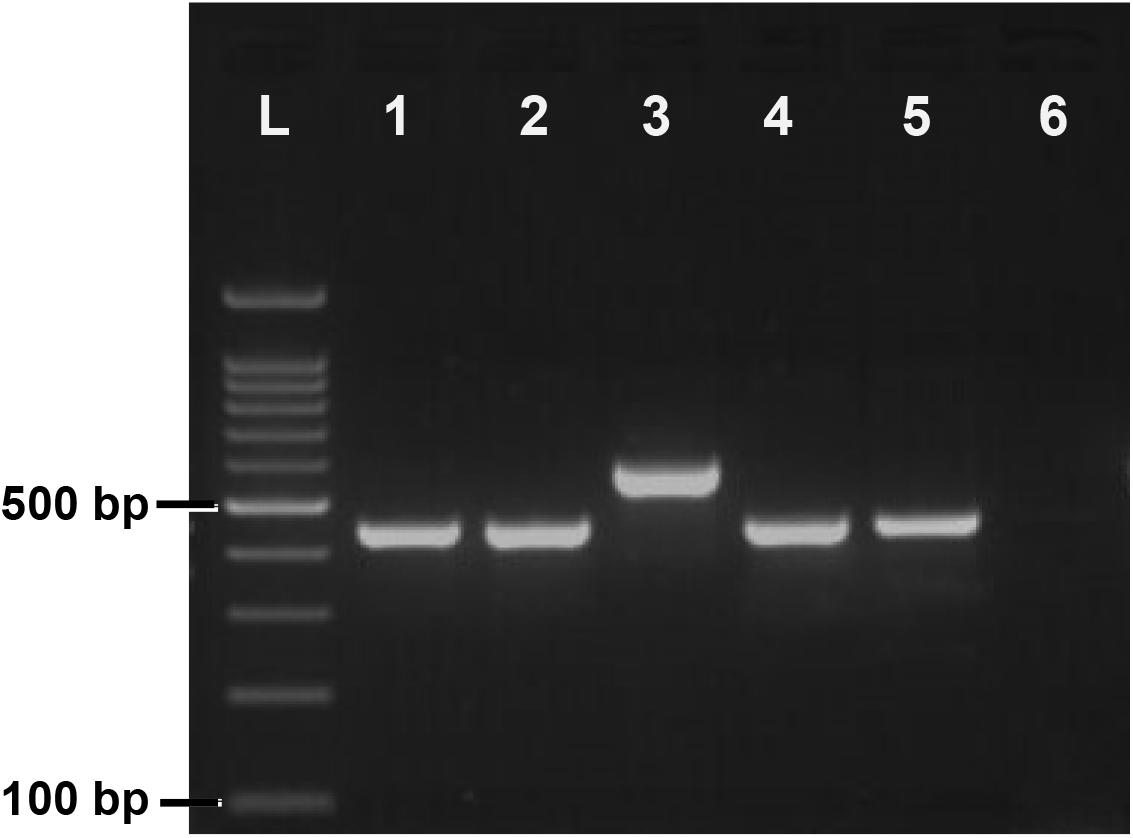
Gel photograph of allele-specific PCR used for the identification of molecular forms A and B in mainland India where only form A and B are prevalent. Lanes L: 100 bp DNA ladder, lane 1: *An. subpictus* Form B, lane 2: *An. subpictus* Form C, lane 3: *An. subpictus* Form A, lane 4: *An. sundaicus* D, lane 5: *An. epiroticus*, lane 6: negative control.

### Distribution of different molecular forms of *An. subpictus s.l.* and *An. sundaicus s.l.*

The distribution of different molecular forms of *An. subpictus s.l.* and *An. sundaicus s.l* in different localities as determined by DNA sequencing as well as by AS-PCR has been shown in **Table 2** and **Figure 1**. It was observed that *An. subpictus* Form A is prevalent throughout mainland India (found exclusively in inland areas) and Sri Lanka, whereas Form B was prevalent only in coastal areas or A&N islands. *Anopheles subpictus* Form C is prevalent only in the A&N Islands. *Anopheles sundaicus* D and *An. epiroticus* were found in Car Nicobar of A&N Islands.

### Assignment of sibling species based on the mode number of float-ridges on eggs

The designation of sibling species of *An. subpictus* collected from mainland India was assigned based on the mode number of ridges on their egg’s float following Suguna et al. (1994) [18]. The distribution of sibling species based on the number of ridges and their molecular forms have been provided in **Table 5**. It was observed that all individuals (n=51) belonging to molecular Form B were correctly assigned to species B based on the mode number of float-ridges. All molecular Form A (n= 97) which were identified as species A, C or D based on the mode number of float-ridges, of which majority (80%) fall under the species C category (**Figure 8**). The mode number of float-ridges on eggs for form B ranged between 15 and 19, and for Form A between 22 and 33 (**Figure 9a**). The number of float-ridges on individual eggs for Form B ranged between 14 and 20, and for Form A ranged between 21 and 35 (**Figure 9b**).

**Table 5.**
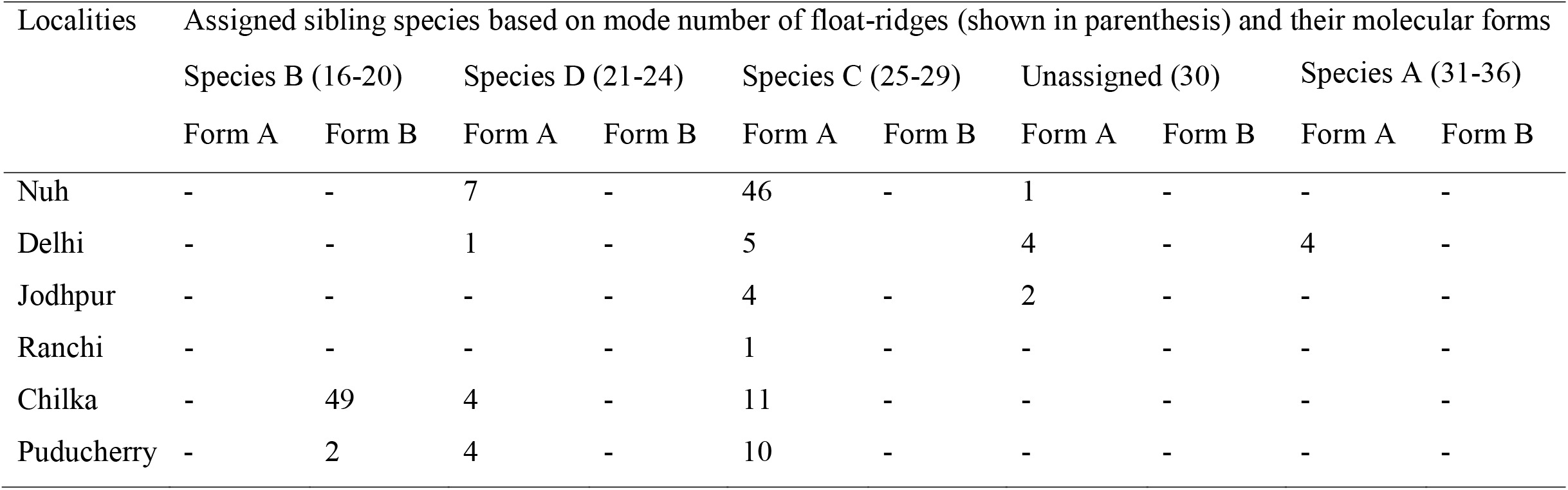
Distribution of *An. subpictus* sibling species based on number of float-ridges on eggs and their molecular forms

| Localities | Assigned sibling species based on mode number of float-ridges (shown in parenthesis) and their molecular forms |  |  |  |  |  |  |  |  |  |
| --- | --- | --- | --- | --- | --- | --- | --- | --- | --- | --- |
|  | Species B (16-20) |  | Species D (21-24) |  | Species C (25-29) |  | Unassigned (30) |  | Species A (31-36) |  |
|  | Form A | Form B | Form A | Form B | Form A | Form B | Form A | Form B | Form A | Form B |
| Nuh | - | - | 7 | - | 46 | - | 1 | - | - | - |
| Delhi | - | - | 1 | - | 5 | - | 4 | - | 4 | - |
| Jodhpur | - | - | - | - | 4 | - | 2 | - | - | - |
| Ranchi | - | - | - | - | 1 | - | - | - | - | - |
| Chilka | - | 49 | 4 | - | 11 | - | - | - | - | - |
| Puducherry | - | 2 | 4 | - | 10 | - | - | - | - | - |

**Figure 8.**
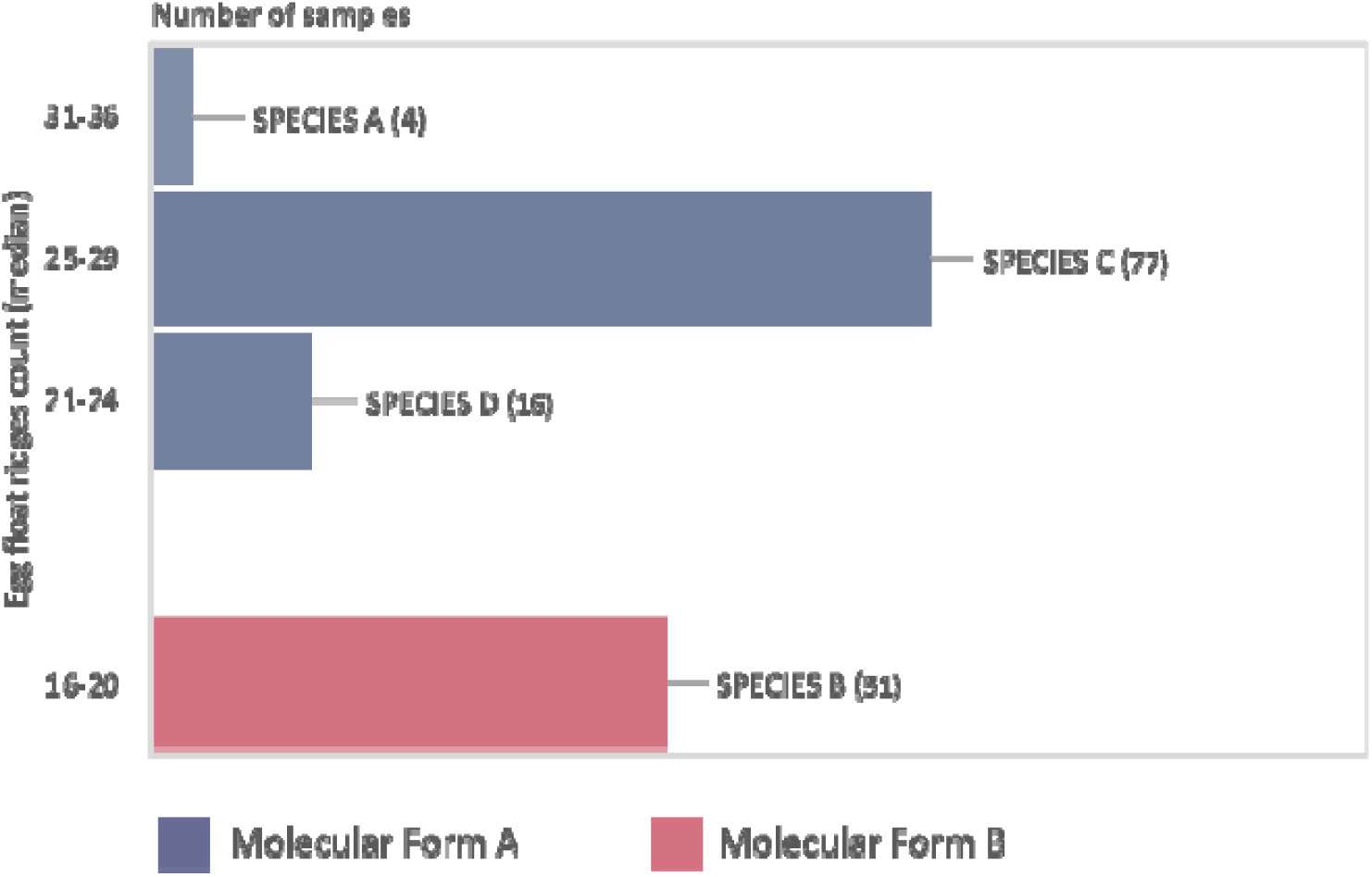
Classification of samples collected from mainland India as sibling species based on mode number of float-ridges [18] and their molecular forms. Details of locations and number of samples have been provided in Table 5.

**Figure 9.**
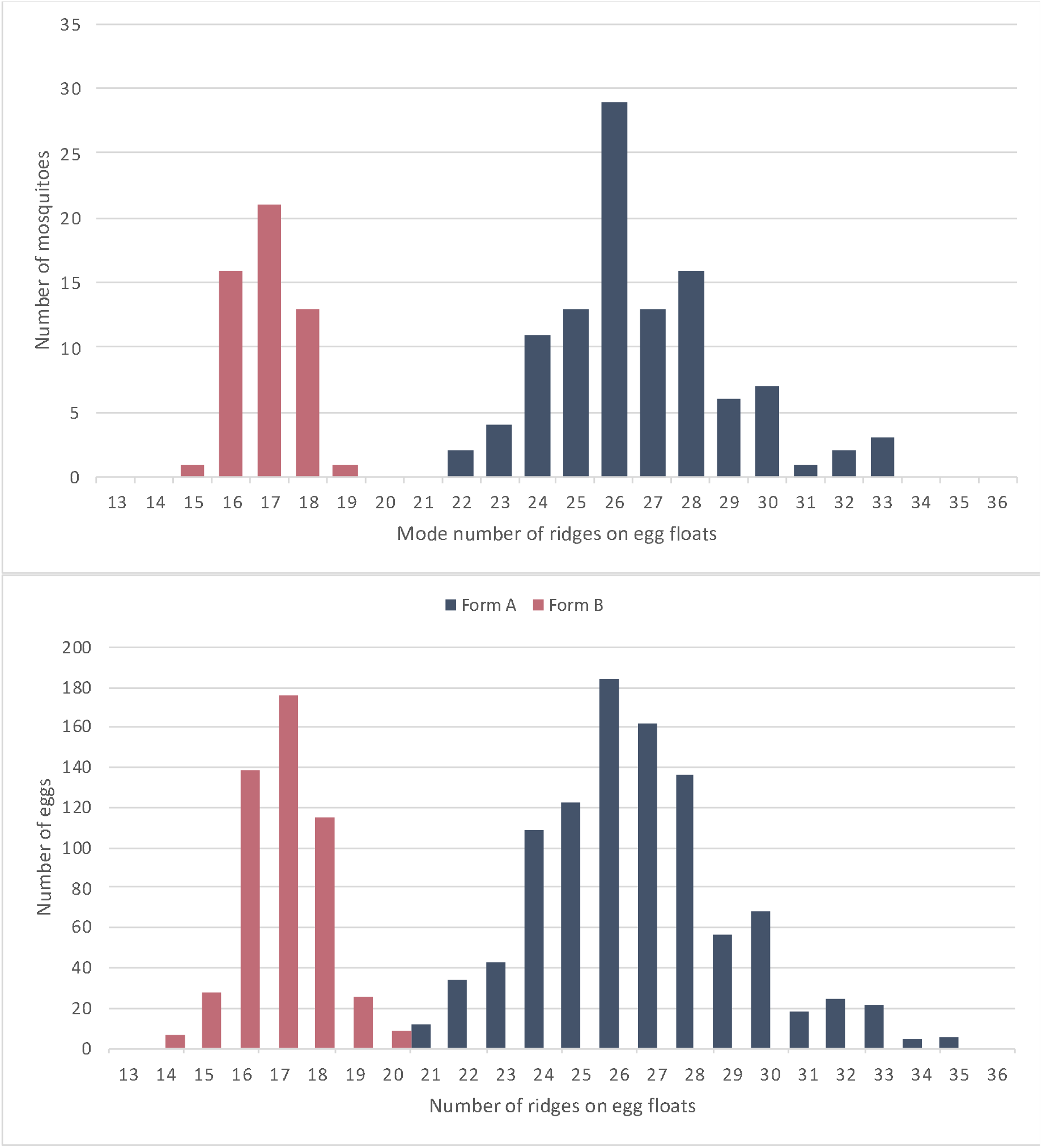
The frequency distribution of mosquitoes with mode number of floats-ridges (a) and eggs with number of float-ridges (b), among molecular form A and B prevalent in mainland India

The frequency distribution of mode number of float-ridges from individual mosquito progeny and number of float-ridges counts from individual egg (pooled data) for molecular Form A and B are shown in **Figure 9**a and b respectively. The figures clearly depict two independent curves, each showing normal distribution, one corresponding to molecular Form B (species B) and another for Form A (comprising of species A, C and D). Based on the distribution pattern of the number of floats-ridges that correlate with molecular forms, it is suspected that species A, C and D might be conspecific while species B is a distinct species.

### Seasonal prevalence of *An. subpictus* Form A and B in areas surrounding brackish water lagoon and near coastal area

A survey was carried out in areas surrounding Chilka lake (a brackish water lagoon) and Puducherry (a coastal area) in monsoon and post-monsoon season, to know the relative prevalence of *An. subpictus* Form A and B in monsoon and post-monsoon season. In the Chilka lake area, only *An. subpictus* Form B was found in the dry season (post-monsoon) when there is no freshwater breeding source available except a large brackish water lagoon. In the monsoon season, when fresh water breeding habitat is prevalent beside brackish water lagoon, Form A was also present (4.7 to 7.6%) alongside Form B. In another area, Puducherry (a coastal area), the proportion of species B was much higher in post-monsoon as compared to autumn. The relative proportion of *An. subpictus* Form A and B in the area surrounding Chilka Lake and coastal area of Puducherry in monsoon and post-monsoon season has given in **Table 6**. The data indicate that the prevalence of Form A of *An. subpictus* determined by the availability of freshwater breeding sources.

**Table 6.**
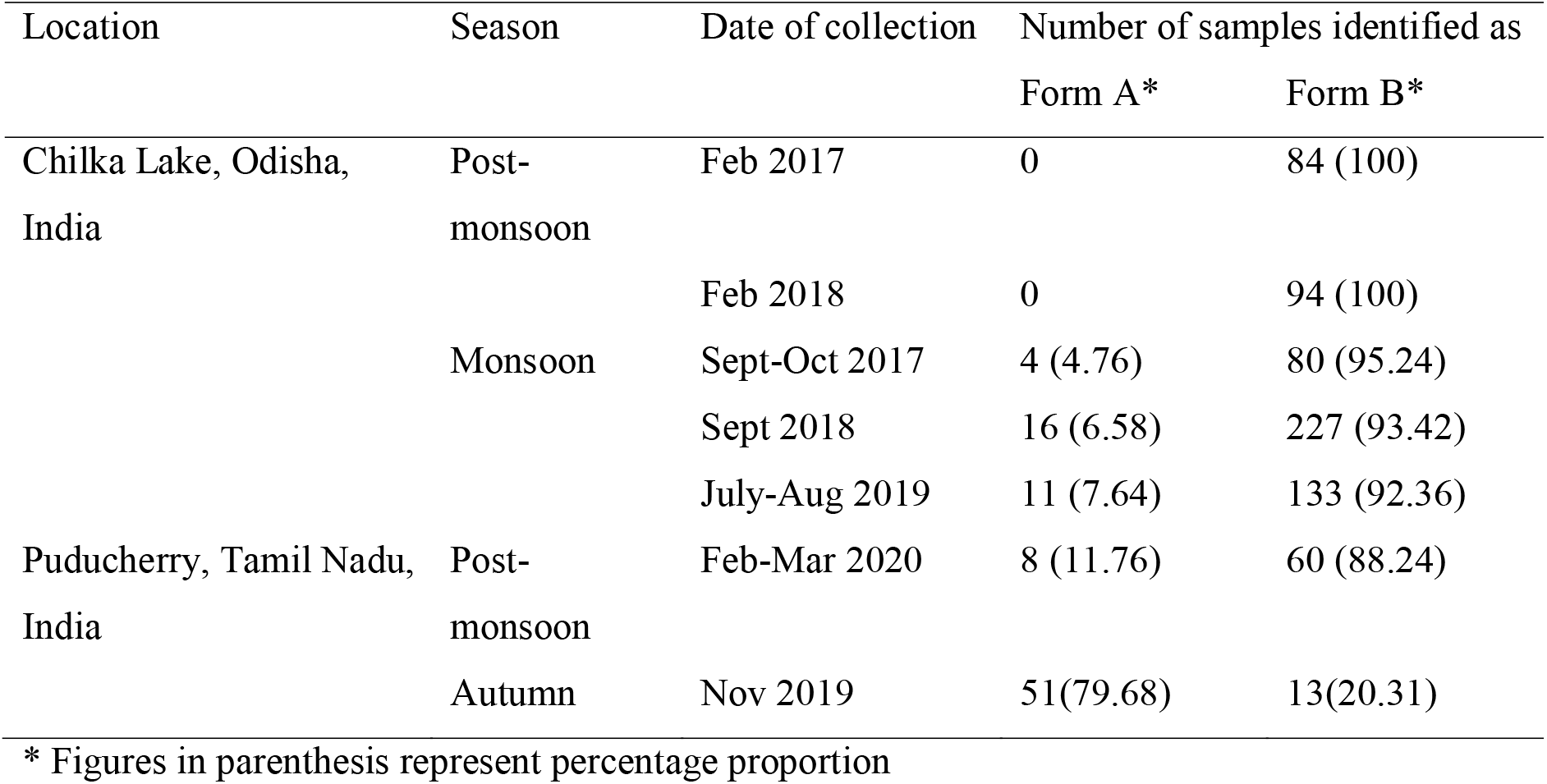
Relative prevalence of *An. subpictus* Form A and B in monsoon and post-monsoon season in areas surrounding Chilka Lake (brackish water lagoon) and Puducherry (coastal area).

| Location | Season | Date of collection | Number of samples identified as |  |
| --- | --- | --- | --- | --- |
|  |  |  | Form A* | Form B* |
| Chilka Lake, Odisha, India | Post-monsoon | Feb 2017 | 0 | 84 (100) |
|  |  | Feb 2018 | 0 | 94 (100) |
|  | Monsoon | Sept-Oct 2017 | 4 (4.76) | 80 (95.24) |
|  |  | Sept 2018 | 16 (6.58) | 227 (93.42) |
|  |  | July-Aug 2019 | 11 (7.64) | 133 (92.36) |
| Puducherry, Tamil Nadu, India | Post-monsoon | Feb-Mar 2020 | 8 (11.76) | 60 (88.24) |
|  | Autumn | Nov 2019 | 51(79.68) | 13(20.31) |

\* Figures in parenthesis represent percentage proportion

## Discussion

Correct identification of anophelines and their sibling species is crucial for the success of any malaria control programme because of possible inherent variations in the epidemiologically-important biological characteristics among sibling species, e.g., vectorial competence [46–47], insecticide resistance [48–51] and host preference [52–53]. In India, *An. sundaicus* is considered as an efficient malaria vector whereas *An.subpictus s.l.* has not been recognized so far as a malaria vector by the national malaria control programme [10]. However, published evidences suggest that at least coastal form of *An. subpictus* is a malaria vector [15, 16]. On the other hand, *An. subpictus* is considered as an important malaria vector in coastal Malaysia [14] and Indonesia [12, 13]. These geographically isolated populations differ in breeding preference, where the Malayan population prefers to breed in brackish water, unlike the majority of Indian *An. subpictus* populations that prefer to breeds in freshwater [14]. In view of the presence of such difference in biological characteristics, recognition of sibling species or ‘fixed’ molecular forms in geographically widely distributed *An. subpictus s.l.* population is an important consideration in understanding their relative epidemiological significance.

So far, four sibling species have been described from a south Indian population of *An. subpictus s.l.* (provisionally designated as species A, B, C and D) based on the arrangement of inversions present on two loci on the polytene chromosome X [18]. To the best of our knowledge, this technique has not been effectively used for the identification of sibling species in *An. subpictus s.l.*, except in a neighbouring country, Sri Lanka [20], where inversions on a single locus (X^+a^ and X^a^), out of two diagnostic loci (X^a/+a^ and X^b/+b^), were examined (which preclude the identification of species C and D). Identification of sibling species in malaria control programme is challenging due to the non-availability of simpler technique, as an alternative to the cytological technique that requires a highly specialized skilled personnel to read banding patterns. Moreover, the cytological technique can be applied only on a fraction of a population, i.e., only live semi-gravid female mosquitoes. As an alternative, molecular tools can be used for the identification of sibling species which are easier and can be applied to dead or alive mosquitoes of either sex. Identification of ‘fixed’ molecular form of rDNA, particularly in ITS2, is a primary step toward the recognition of sibling species. In this direction, ITS2 of *An. subpictus s.l.* populations have been characterized in Sri Lankan [27], Thai and Indonesian population [30], however, reliable data on molecular forms from India is lacking. The present study is an attempt to characterize all the molecular forms of morphologically identified *An. subpictus s.l.* and *An. sundaicus s.l.* prevalent in the Indian subcontinent to uncover hidden cryptic species, distribution pattern and to establish their phylogenetic relationship. Characterization of molecular forms will help in identifying sibling species and developing a reliable and simpler method for their identification. Characterization of *An. sundaicus s.l.* was included in this study in view of the report of incongruence in molecular taxonomy with formal (morphological) taxonomy, where one molecular form of *An. subpictus s.l.* (mostly species B) was found to be a close relative of *An. sundaicus s.l* [27]. As a result, Surendran et al. (2010) [27] recognized them as a member of the Sundaicus Complex. Earlier phylogenetic study on Sri Lankan *An. subpictus* [27, 28] was based on partial ITS2 and D3 domain of 28S rDNA. Here we used full length ITS2 sequence and 28S-D1-D3 to identify molecular forms of both *An. subpictus s.l.* and *An. sundaicus s.l.* from the Indian subcontinent.

This study revealed the presence of a total of five molecular forms in Sundaicus-Subpictus complex in the Indian subcontinent (both by ITS2 as well as 28S-D1-D3): three molecular forms present in morphologically identified *An. subpictus* (Form A, B and C) and two molecular forms in *An. sundaicus* complex (*An. epiroticus* and *An. sundaicus* D). Phylogenetic analysis using two regions of rDNA revealed two distinct clades where *An. subpictus* Form B and C, *An. epiroticus, An. sundaicus* and *An. indefinitus* falls under one clade (Sundaicus) and one molecular form, Form A, falls under distantly related clade (Subpictus). Thus, the phylogenetic relationship showed incongruence with the morphological delimitation of the species. In an earlier study, Surendran et al. (2010) [27] showed that most of the *An. subpictus* species B (Form B in our study) from Sri Lanka fall under Sundaicus complex and therefore considered as a member of Sundaicus Complex. We include another molecular form of *An. subpictus*, i.e., Form C, found in A&N Island as a member of Sundaicus Complex. The two molecular forms very recently reported from Indonesia and Thailand [30] also fall in the same clade and closely related to Indian molecular forms Form B and C.

The designation of molecular forms in this study is based on ribosomal DNA (ITS2 and 28SrDNA), where we didn’t find any intraspecific variation in a specific population (due to concerted evolution), as seen in mitochondrial DNA (maternally inherited and mutations are passed through generations). The absence of individual variation in DNA sequences in molecular forms of *An. subpictus* across wide geographical areas is contrary to the findings by Kaura et al. (2010) [31], Chattopadhyay et al. (2013) [32] and Chhilar et al. [33], where they recorded individual variation in ITS2 sequences in samples collected from inland India. These sequences are very similar to Form A and we believe that individual differences in ITS2 sequences presented in above-mentioned publications is due to errors in the sequence-read of Form A arising due to the 12 bp indel in one haplotype leading to ambiguous sequence beyond the point of indel in sequence read. Interestingly, the intragenomic variation in Form A, with the presence of two haplotypes, was found fixed throughout its distribution, from north India to Sri Lanka. The fixed molecular forms based on rDNA, which are found in sympatricity, deserve distinct specific status, but the specific status of closely related molecular forms, which are allopatric, is obscure. Thus the two molecular forms A and B found in sympatry in southern Asia (India and Sri Lanka) are genetically isolated but the specific status of other molecular forms found in South-east Asia (A&N Islands, Thailand and Indonesia) is not absolute. Among Sundaicus Complex, *An. sundaicus* D and *An. epiroticus* have been found sympatric in this study in Car Nicobar Island, and they may be considered distinct biological species.

Wilai et al. [30] reported that *An. subpictus s.l.* from Thailand and Indonesia are different from India and Sri Lanka. They recorded two types of ITS2 sequences from Thailand which were identical except in one type of sequence. There was a mixed nucleotide base at one position. It is unlikely that these two forms are reproductively isolated. The specific status of allopatric molecular forms cannot be ascertained using population genetic analysis, which can be verified using crossing experiments only. Recently, Wilai et al. [30] has carried out crossing experiments in two forms having different mtDNA sequence (but same molecular forms with respect to ITS2) and found genetic compatibility between them. We consider these two mtDNA forms as intraspecific variation because such variation is common in mtDNA being inherited maternally. Such intraspecific variation were also found in this study where we recorded multiple sequence of *COII* mtDNA, even in small numbers of samples. The molecular forms reported from Thailand and Indonesia were very close to Indian molecular forms B and C with a very low genetic distance ranging from 0.0 to 0.018 (with respect to ITS2). The lack of genetic distance between Indonesian *An. subpictus* and Form C was due to 100% similarity in sequences except presence of two bp indel in Form C.

Interestingly two molecular forms of *An. subpictus s.l.*, Form B and C, which falls under sundaicus clade, are found in coastal areas or islands. This may be due to their breeding preference is brackish water, similar to most of the members of *An. sundaicus* complex. This is complemented by the fact that, in Chilka Lake (a brackish lagoon), we found exclusively Form B in Post monsoon season (October and February), while this was present along with Form A in Monsoon season (August-September) when freshwater breeding habitats are abundantly present beside brackish water habitat. Suguna (1982) [29] and Panicker et al. (1981) [15] also found species B of Subpictus Complex along with species A in the rainy season. These observations indicate that Form B and Form C of *An. subpictus*, found in coastal areas or islands, has a preference to breed in brackish water. Another form of *An. subpictus*, Form A, which is widely distributed throughout inland India, appears to breed in freshwater.

Review of literature shows that species B of *An. subpictus*, found mainly in coastal areas [19, 20], is a potential malaria vector [15, 54]. Recently *An. subpictus s.l.* was reported be a potential malaria vector from Goa (coastal area) [16]. This study confirms presence of molecular form B in Goa, It appears that molecular Form B (species B) is playing role in malaria transmission, however, further study is warranted. However, in some inland areas, *An. subpictus* has been found to have sporozoite in their salivary glands [54, 55, 56]. The differential role of these molecular forms in malaria transmission need to be confirmed.

We attempted to classify sibling species of Form A and B of the Subpictus Complex, prevalent in mainland India based on the mode number of float-ridges. All the molecular Form B were correctly classified as species B while the majority of Form A were characterized as species C. Frequency distribution pattern of float-ridges clearly shows two distinct peaks, one peak corresponds to molecular Form A and another peak corresponds to Form B. It appears that all molecular Form A is a single species. Because majority of species C identified based on ridges on egg-floats belong to molecular Form A, it is erroneous to assign them as ‘Species A’ [27, 57]. It is recommended that they may be described as molecular Form A instead of Species A. However, molecular Form B may be designated as ‘Species B’. It is evident from our data that molecular method as well as mode number of float ridges can correctly identify species B, and can differentiate them from molecular Form A. However, morphological identification based on egg’s ridges has certain limitations. It cannot differentiate species B from *An. sundaicus s.l.* due to overlapping range of ridges (range 16 to 20). Moreover, morphological identification based on the number of ridges is a cumbersome process and can be used with only live and gravid female mosquitoes, whereas molecular method can be applied on dead or alive mosquitoes irrespective of life stages and sex.

*Anopheles sundaicus s.l.* is widely distributed species and a malaria vector in the coastal region of Southeast Asia [25]. In India, this is the most prevalent and important malaria vector in the A&N Islands. Earlier studies by Nanda et al. (2004) [22] and Alam et al. (2005) [23] revealed the presence of only one sibling species in A&N Islands, i.e., *An. sundaicus* D (variant III). This study confirms the presence of *An. sundaicus* D in A&N islands (Port Blair and Car Nicobar). We also recorded the presence of *An. epiroticus* in Car Nicobar island for the first time, though just one specimen out of small sample size (n=52). Further studies are required to explore the presence of this species or new cryptic species in this area. In addition, we recorded the presence of *An. epiroticus* in Myanmar, which is in agreement with Dusfour et al. (2007) [24]. However, *An. epiroticus* recorded from Car Nicobar island was having mixed haplotypes of ITS2, unlike *An. epiroticus* from Myanmar or classic *An. epiroticus*. A similar variant has been reported in Indonesia, Thailand and Vietnam [58, 59]. Dusfour et al. (2007) [24] reported combinations of variants I and III, II and III, and I, II and III in continental Southeast Asia. Syafruddin et al. (2020) [59] hypothesized that such mixed bases may be due to possible hybridization/introgression between sympatric sibling species.

In this study, a new PCR based assay was developed for the differentiation of members of sundaicus and subpictus clade. Initially, it was designed to differentiate Form A and B of Subpictus Complex but, later upon discovery of Form C, we found that this could not differentiate Form B and C. However, this PCR can differentiate unambiguously members of Sundaicus (*An. subpictus* Form A) and subpictus clade (*An. subpictus* Form B, Form C and *An. sundaicus s.l.*). A similar ASPCR assay was developed by Surendran et al. (2013) [28], but we developed a new assay because we noticed that this ASPCR has cross-reactivity with other Indian malaria vector *An. stephensi*, which has superficial resemblance with *An. sundaicus s.l.* [26] and often misidentified in old specimens with lost palpal ornamentation and scales.

This study has certain limitation. As we do not have data on inversion genotypes of the polytene chromosome of *An. subpictus s.l.*, we could not correlate association of molecular forms with previously designated sibling species. Recently one new species *An. pseudosundaicus* belonging to the Pyretophorus series [60] has been designated from Kerala, southwest India, which unlike *An. sundaicus s.l.* doesn’t have speckling on legs and closer to *An. subpictus* based on the DNA sequence of *cytochrome oxidase C subunit 1*. However, access to this molecular data has not been made public. Therefore, it could not be ascertained if this species is one of the molecular forms we described in this study. A more elaborate study on molecular characterization of all cytological forms of *An. subpictus* is desirable.

## Conclusions

Based on nuclear rDNA sequences, we identified three fixed molecular forms (Form A, B and C) in the morphologically classified *An. subpictus s.l.* and two forms in *An. sundaicus s.l.* in the Indian subcontinent. Molecular phylogenetic analysis revealed two diverse clades of mosquitoes where two molecular forms (B and C) of morphologically identified *An. subpictus* found mainly in coastal areas or islands along with molecular forms reported from Thailand and Indonesia were found to be close relative of *An. sundaicus s.l.* A molecular form of *An. subpictus*, Form A, prevalent in mainland India and Sri Lanka, has fixed intragenomic sequence variation in ITS2 throughout its distribution with the presence of two haplotypes differing by 12 bp indel. The presence of *An. epiroticus* was recorded first time in Car Nicobar (A&N Islands, India).

## Additional files

**Additional file 1: Figure S1** Alignment of rDNA sequences (ITS2 to D3 domain of 28S) of different molecular forms of *An. subpictus s.l.*, and *An. sundaicus* complex *s.l.*.

## Abbreviations

A&N: Andaman & Nicobar
ASPCR: 
bp: base pair
*COI*: *cytochrome oxidase I*
*COII*: *cytochrome oxidase II*
ITS2: Internal Transcribed Spacer-2
min: minute
PCR: Polymerase Chain Reaction
rDNA: ribosomal DNA
s: second
TE: Tris-EDTA

## Acknowledgements

Authors are thankful to Dr U Pe Htun, Medical Entomology Research Division, Yangon, Myanmar for providing *An. sundaicus s.l.* samples. Thanks are due to Mr Uday Prakash, Mr NS Bhakuni and Mrs Alka Kapoor for processing of samples and performing PCR reactions; Mr Kanwar Singh, Mr Bhupal Ram, Mr Sadruddin for sample collections.

## Ethics approval and consent to participate

Not applicable.

## Consent for publication

Not applicable.

## Availability of data and materials

All data generated or analyzed during this study are included in this published article and its additional files.

## Competing interests

The authors declare that they have no competing interests.

## Funding

Financial support to this study was provided by the Department of Science and Technology (DST), Government of India, under Women Scientist Scheme-B (Grant no. SR/WOS-B/698/2016).

## Authors’ contributions

OPS conceived and designed study, designed PCR and sequencing strategies; AS performed PCR, DNA sequencing, phylogenetic analysis; GS performed cloning experiments; MKD and SNS supervised mosquito sampling from field and provided critical suggestions to the manuscript; BRK and HPL supervised the study and contributed to the manuscript. OPS and AS wrote the first draft of the manuscript and all authors provided critical input to the manuscript and approved the final manuscript

## Notes

### Competing Interest Statement

The authors have declared no competing interest.

