## Supplementary material for "Molecular forms of *Anopheles subpictus s.l.* and *Anopheles sundaicus s.l.* in the Indian subcontinent": Figure S1

**Additional file 1:**

**Figure S1.** Alignment of rDNA sequences (ITS2 to D3 domain of 28S) of different molecular forms of *An. subpictus* and *An. sundaicus* complex. Sub_A_h1= *An. subpictus* Form A (haplotype 1), Sub_A_h2= *An. subpictus* Form A (haplotype 2), Sub_B= *An. subpictus* Form B, Sub_C= *An. subpictus* Form C, Epi_I = *An. epiroticus* (variant I) from Myanmar, Epi_II = *An. epiroticus* from Car Nicobar, Sun_D = *An. sundaicus* D (variant III). MT068436, MT068425 and MT068434 are representative sequences of *An. subpictus* from Thailand and Indonesia [1]. Dots in alignment represent similarity in nucleotide base with the first row of sequence and dash represents missing nucleotide. The demarcation of ITS2-boundries was done using online ITS2 database tool available at http://its2.bioapps.biozentrum.uni-wuerzburg.de/ (Selig et al., 2008) [2]

```
         ──────────────────────────────────────5.8S─────────────><────────ITS2───────────
Sub_A_h1 CGCATATGGCGCATCGGACGTTTCAACCCGACCGATGCACACATCCTTGAGTGCCTACTAGGTACTGAGAGATTCCTATA [80]
Sub_A_h2 ................................................................................
Sub_B    ..................................................................TC..TT........
Sub_C    ..................................................................TC..TT........
Epi_I    ..................................................................TC..TT........
Epi_II   ..................................................................TC..TT........
Sun_D    ..................................................................TC..TT........
MT068436 ..................................................................TC..TT........
MT068425 ..................................................................TC..TT........
MT068434 ...........Y......................................................TC..TT........
         ──────────────────────────────────────ITS2──────────────────────────────────────
Sub_A_h1 ACTTGACTACAGACGGGCGCCACAAACGGGCTGACGGGCCATCCGTCGTCCGGCGTGCGACTGTGCAGCATGGCGTGCTC [160]
Sub_A_h2 ................................................................................
Sub_B    .T.A...................T..T...........TT..........T.............................
Sub_C    CT.A...................T..............TT..........T.............................
Epi_I    .T.A...................T...............T..........T.............................
Epi_II   .T.A...................T...............T..........T.............................
Sun_D    .T.A...................T...............T..........T.............................
MT068436 CT.A...................T..............TT..........T.............................
MT068425 .T.A...................T..............TT..........T.............................
MT068434 .T.A...................T..............TT..........T.............................
         ──────────────────────────────────────ITS2──────────────────────────────────────
Sub_A_h1 GGGTCTCGGCGTGGACCCTTGGGCGCTGAAAGTGGACACT--GTTTG-GCGGCACCTGCGCGTGTGCTCTC---AGTGTT [240]
Sub_A_h2 ........................................--.....-.......................---......
Sub_B    ....................................T...CT.....A.........TT............CTA.....C
Sub_C    ....................................T...CT.....A.........TT............CTA.....C
Epi_I    ....................................T...CT.....A.........TT............CTA.....C
Epi_II   ....................................T...CT.....A.........TT............CTA.....C
Sun_D    ....................................T...CT.....A.........TT............CTA.....C
MT068436 ....................................T...CT.....A.........TT............CTA.....C
MT068425 ....................................T...CT.....A.........TT............CTA.....C
MT068434 ....................................T...CT.....A.........TT............CTA.....C
         ──────────────────────────────────────ITS2──────────────────────────────────────
Sub_A_h1 GACGTATGGTGAGGGTAGTGTCAAATCGCACGGTTCGACAACA--AGCGTACCGTCGAGTTTGGTGCAATCGGATGCCTA [320]
Sub_A_h2 ...........................................--...................................
Sub_B    ........................GC........G........CA.......T...........................
Sub_C    ........................GC........G........CA.......T...........................
Epi_I    ........................GC........G........CA.......T...........................
Epi_II   ........................GC........G........CA.......T...........................
Sun_D    ........................GC........G........CA.......T...........................
MT068436 ........................GC........G........CA.......T...........................
MT068425 ........................GC........G........CA.......T...........................
MT068434 ........................GC........G........CA.......T...........................

         ──────────────────────────────────────ITS2──────────────────────────────────────
Sub_A_h1 CTACCATGGGCGGAGCCGGCGTGCATTCAACACTCGACGT------CCTGTATCAACCGGATGCCAACTTGGTTGGTGGT [400]
Sub_A_h2 ........................................------..................................
Sub_B    .............T..........................GTGCGT......................------------
Sub_C    .............T.............A............GCGTGT.............R........------------
Epi_I    .............T..........................GCGTGT......................------------
Epi_II   .............T..........................GCGTGT......................------------
Sun_D    .............T..........................GCGTGT......................------------
MT068436 .............T.............A............GCGTGT......................------------
MT068425 .............T..........................GTGTGT......................------------
MT068434 .............T..........................GTGTGT......................------------
```

```
            ————————————————————————————————————ITS2————————————————————————————————————
Sub_A_h1    GCCGGCGCAGACAGGACACTGAATCGATCTTGGTGGTACAACCCACATGTGGGTTAGTAGGTAGGTGGTCGATGTGTGCA [480]
Sub_A_h2    ..................................................------------..................
Sub_B       --------------------------------------------------------------------------------
Sub_C       --------------------------------------------------------------------------------
Epi_I       --------------------------------------------------------------------------------
Epi_II      --------------------------------------------------------------------------------
Sun_D       --------------------------------------------------------------------------------
MT068436    --------------------------------------------------------------------------------
MT068425    --------------------------------------------------------------------------------
MT068434    --------------------------------------------------------------------------------

            ————————————————————————————————————ITS2————————————————————————————————————
Sub_A_h1    GGTGACAACCGGATGCCAGCGATGGCGGTGCCGGCGCACACCAGCACACTCGCCCCTAGGTCGCTTGTTGCGTGTAACGC [560]
Sub_A_h2    ................................................................................
Sub_B       ----------T.C..--TCA.T...T....T.......G..AG.---..G...G.G..---......AGT..........
Sub_C       ----------T.C..--TCA.T...T....T.......G..AG.---..G...G.G..---......AGT..........
Epi_I       ----------T.C..TGTCA.T...T....T.......G..AG.---..G...G.G..---......AGT..........
Epi_II      ----------T.C..TGTCA.T...T....T.......G..AG.---..G...G.G..---......AGT.......K..
Sun_D       ----------T.CC.TGTCA.T...T....T.......G..AG.---..G...G.G..---......AGT..........
MT068436    ----------T.C..TGTCA.T...T....T.......G..AG.---..G...G.G..---......AGT..........
MT068425    ----------T.C..--TCA.T...T....T.......G..AG.---..G...G.G..---......AGT..........
MT068434    ----------T.C..--TCA.T...T....T.......G..AG.---..G...G.G..---......AGT..........

            ————————————————————————————————————ITS2————————————————————————————————————
Sub_A_h1    GTGTGATCCATACACATACCTGTTTGAGCTGTGCGTTGAACACAAGAGGATGAGAGTTGCCAAACA---CACAGACCACA [640]
Sub_A_h2    ..................................................................---...........
Sub_B       ...C..C........G......C.......A.......------C...TTG...........G..GGAGG.A.T......
Sub_C       ...C..C........G......C.......A.......------C...TTG...........G..GGAGG.G.T......
Epi_I       ...C..C........G......C...............------C...CTG...........G..GGAG..A.T......
Epi_II      ...C..C........G......C...............------C...CTG...........G..GGAG..A.T......
Sun_D       ...C..C........G......C...............------C...CTG...........G..GGAG..A.T......
MT068436    ...C..C........G......C.......A.......------C...TTG...........G..GGAGG.G.T......
MT068425    ...C..C........G......C.......A.......------C...TTG...........G..GGAG..A.T......
MT068434    ...C..C........G......C.......A.......------C...TTG...........G..GGAG..A.T......

            ————ITS2———><———————————————————————28S—————————————————————————————————————
Sub_A_h1    ----CTCCAGTAGGCCTCAAGTGATGTGTGACTACCCCCAGAATTTAAGCATATTAATAAGGGGAGGAAGAGAAACCAAC [720]
Sub_A_h2    ----............................................................................
Sub_B       ACTA............................................................................
Sub_C       AACT............................................................................
Epi_I       --TA............................................................................
Epi_II      --TA............................................................................
Sun_D       --TA............................................................................
MT068436    AACT.............................
MT068425    CCTA.............................
MT068434    CCTA.............................

            ————————————————————————————————————28S—————————————————————————————————————
Sub_A_h1    CGGGATTCCCTGAGTAGCTGCGAGCGAAACGGGAGAAGCCCAGCACGTAGGGACGACGTGGAGACGCGCCTGTCCGGTTC [800]
Sub_A_h2    ................................................................................
Sub_B       .......................................T...............G.........A..............
Sub_C       .......................................T...............G........................
Epi_I       .......................................T...............G........................
Epi_II      .......................................T...............G........................
Sun_D       .......................................T...............G........................

            ————————————————————————————————————28S—————————————————————————————————————
Sub_A_h1    CGTGTATTGGACCGGTCCGTTATCTATCATGTACAGTGTGCATCAAGTCTAACTTGAAGGTGGCTCATTATCCCATAGAG [880]
Sub_A_h2    ................................................................................
Sub_B       ...............................C......CA........................................
Sub_C       ...............................C......CA........................................
Epi_I       ...............................C......CA........................................
Epi_II      ...............................C......CA........................................
Sun_D       ...............................C......CA........................................

            ————————————————————————————————————28S—————————————————————————————————————
Sub_A_h1    GGTGATAGGCCCGTTGCATGCGCGCGCCTGTGCTGGTAGACGGTCGGCTCCAGGGAGTCGTGTTGCTTGATAGTGCAGCA [960]
Sub_A_h2    ................................................................................
Sub_B       ..............A........A........................................................
Sub_C       ..............A........A........................................................
Epi_I       ..............A........A........................................................
Epi_II      ..............A........A........................................................
Sun_D       ..............A........A........................................................
```

```
          ─────────────────────────────────────28S──────────────────────────────────────
Sub_A_h1 CTAAGTGGGAGGTAAACTCCTTCTAAGGCTAAATATGACCATGAGACCGATAGCGAACAAGTACCGTGAGGGAAAGTTGA [1040]
Sub_A_h2 ................................................................................
Sub_B    ..........................A.........CG..........................................
Sub_C    ..........................A.........TG..........................................
Epi_I    ..........................A.........TG..........................................
Epi_II   ..........................A.........TG..........................................
Sun_D    ..........................A.........TG..........................................

          ─────────────────────────────────────28S──────────────────────────────────────
Sub_A_h1 AAAGCACTCTGAATAGAGAGTCAAATAGTACGTGAAACTGCCTAGGGTACACAAACCCGTTGAACTCAATGATCCGGGCG [1120]
Sub_A_h2 ................................................................................
Sub_B    ................................................................................
Sub_C    ................................................................................
Epi_I    ................................................................................
Epi_II   ................................................................................
Sun_D    ................................................................................

          ─────────────────────────────────────28S──────────────────────────────────────
Sub_A_h1 GCGATATTCAGCGGCGGTCGCAAGGCCACCGTGCACTTATCGTTCCGCAGCAAACGGACATCGCGATCCATTACAATAGC [1200]
Sub_A_h2 ................................................................................
Sub_B    ..............T............G....................................................
Sub_C    ..............T............G....................................................
Epi_I    ..............T............G....................................................
Epi_II   ..............T............G....................................................
Sun_D    ..............T............G....................................................

          ─────────────────────────────────────28S──────────────────────────────────────
Sub_A_h1 GAGTCGAGCGTTCTGCGCTGCGACCGTCCGGCAT-TACTGGCCCCTGGCTCGTGGTGGACGGCTCCCTAGTAGGGGCGGC [1280]
Sub_A_h2 ..................................-.............................................
Sub_B    .G....G...C.T.....................G.TA..........................................
Sub_C    .GT...G...C.T.....................GCGA..........................................
Epi_I    .G....G...C.T.....................GCGA..........................................
Epi_II   .G....G...C.T.....................GCGA..........................................
Sun_D    .G....G...C.T.....................GCGA..........................................

          ─────────────────────────────────────28S──────────────────────────────────────
Sub_A_h1 TTGACGGCCGCACCGAGCGGGGGTCTCCGCGCCTTTCTTCTGAAGGGCGACTGGGTCCGACCGAGCGTGGTGTGCCGCTG [1360]
Sub_A_h2 ................................................................................
Sub_B    ..TG.................................A..C.......................................
Sub_C    ..TG.................................A..C.........M.............................
Epi_I    ..TG.................................A..C.......................................
Epi_II   ..TG.................................A..C.......................................
Sun_D    ..TG.................................A..C.......................................

          ─────────────────────────────────────28S──────────────────────────────────────
Sub_A_h1 GAACCGTGATGGATCCGAGCGTGGGGCCGACC--AAGGCTGCCCTC----GTGGCCAGCCAAGCGTGCCCCCAGATCGGC [1440]
Sub_A_h2 ................................--............----..............................
Sub_B    .........................A....GTTAG....A...T..TGCGAA..TTGT..GTT....T............
Sub_C    .........................A....GTTAG....A...T..TGCGAA..TTGT..GTT....T............
Epi_I    .........................A....GTTAG....A...T..TGCGAA..TTGT..GTT....T............
Epi_II   .........................A....GTTAG....A...T..TGCGAA..TTGT..GTT....T............
Sun_D    .........................A....GTTAG....A...T..TGCGAA..TTGT..GTT....T............

          ─────────────────────────────────────28S──────────────────────────────────────
Sub_A_h1 GATGAGCAGCATGCATTGAGGCACCTCCGGGACCCGTCTTGAAACACGGACCAAGAAGTCTATCTTGCACGCGAGCCAAT [1520]
Sub_A_h2 ................................................................................
Sub_B    .....................T..................................................A.......
Sub_C    .....................T..................................................A.......
Epi_I    .....................T..................................................A.......
Epi_II   .....................T..................................................A.......
Sun_D    .....................T..................................................A.......

          ─────────────────────────────────────28S──────────────────────────────────────
Sub_A_h1 GGGACCGGTGCCACCTTCGGGTGGTGCCGCTCAATCGAACACCCATAGGCGAAGACAACTCGAAGACGCATGTCACGGGA [1600]
Sub_A_h2 ................................................................................
Sub_B    .......A.A.....GGT.T......T....................................T....TTC.........
Sub_C    .......A.A.....GGT.T......T....................................T....TTC.........
Epi_I    .......A.A.....GGT.T......T....................................T....TTC.........
Epi_II   .......A.A.....GGT.T......T....................................T....TTC.........
Sun_D    .......A.A.....GGT.T......T....................................T....TTC.........

          ─────────────────────────────────────28S──────────────────────────────────────
Sub_A_h1 TTACGGGTTCGGCATTGGCGCAAGCCTACGTCGGATCCCTCCATCCCAGGGTGCCCCGATACGGGTGGGAGCGGGCCTCC [1680]
Sub_A_h2 ................................................................................
Sub_B    ...............-...........T....................................................
Sub_C    ...............-...........T....................................................
Epi_I    ...............-...........T....................................................
Epi_II   ...............-...........T....................................................
Sun_D    ...............-...........T....................................................
```

```
                    ————————————————————————————————————————28S————————————————————————————————————————
Sub_A_h1 GGGCGTCGTTCTACCCAGCGGGCATACCCCGAGTGTGCAGGATGCGACCCGAAAGATGGTGAACTATGCCTGATCAGGTT [1760]
Sub_A_h2 ................................................................................
Sub_B    ................................................................................
Sub_C    ................................................................................
Epi_I    ................................................................................
Epi_II   ................................................................................
Sun_D    ................................................................................

                    ————————————————————————————————————————28S———————————————
Sub_A_h1 GAAGTCAGGGGAAACCCTGATGGAGGACCGAAGCAATTCTGACGTGCAAATCGAT [1815]
Sub_A_h2 .......................................................
Sub_B    .......................................................
Sub_C    .......................................................
Epi_I    .......................................................
Epi_II   .......................................................
Sun_D    .......................................................
```
